# SSMART: Sequence-structure motif identification for RNA-binding proteins

**DOI:** 10.1101/287953

**Authors:** Alina Munteanu, Neelanjan Mukherjee, Uwe Ohler

## Abstract

**Motivation:** RNA-binding proteins (RBPs) regulate every aspect of RNA metabolism and function. There are hundreds of RBPs encoded in the eukaryotic genomes, and each recognize its RNA targets through a specific mixture of RNA sequence and structure properties. For most RBPs, however, only a primary sequence motif has been determined, while the structure of the binding sites is uncharacterized.

**Results:** We developed **SSMART**, an RNA motif finder that simultaneously models the primary sequence and the structural properties of the RNA targets sites. The sequence-structure motifs are represented as consensus strings over a degenerate alphabet, extending the IUPAC codes for nucleotides to account for secondary structure preferences. Evaluation on synthetic data showed that **SSMART** is able to recover both sequence and structure motifs implanted into 3‘UTR-like sequences, for various degrees of structured/unstructured binding sites. In addition, we successfully used **SSMART** on high-throughput *in vivo* and *in vitro* data, showing that we not only recover the known sequence motif, but also gain insight into the structural preferences of the RBP.

**Availability:** **SSMART** is freely available at https://ohlerlab.mdc-berlin.de/software/SSMART_137/

## 1 Introduction

RNA-binding proteins (RBPs) are key players in RNA metabolism and function. They bind RNA molecules through cis-regulatory elements to coordinate post-transcriptional processes such as splicing, RNA transport, RNA stability and localization (Keene, 2007). There are hundreds of RBPs encoded in eukaryotic genomes (Baltz *et al.*, 2012), each with specific functions, thus it is necessary that the RBPs recognize their RNA targets with high specificity. The current understanding is that this binding specificity is achieved through combinations of RNA sequence and structure properties, in variable proportions (Cook *et al.*, 2014). Some RBPs prefer to bind single-stranded RNA and recognize their target only by the nucleotide composition, while others prefer specific structural contexts. For most RBPs, however, only a primary sequence motif has been determined, while the structure of the binding sites is uncharacterized.

RBP-RNA interactions are experimentally assessed with high-throughput *in vitro* or *in vivo* methods. The *in vitro* approaches, like RNAcompete (Ray *et al.*, 2009), determine the binding specificity and affinity of a specific protein to millions of short, synthetic RNAs, in the absence of other proteins or cellular factors, while the *in vivo* methods ascertain the binding sites of a certain protein in a specific cellular context. There are a number of crosslinking and immuno-precipitation (CLIP) methods (Ule *et al.*, 2003; Konig *et al.*, 2010; Hafner *et al.*, 2010) that induce permanent cross-links between RNAs and RBPs *in vivo*, after which the RBP-RNA fragments are isolated using immunoprecipitation, and the crosslinked RNA segments are sequenced.

Computational analysis of RBP-RNA interactions is vital for interpreting the experimental data and finally understanding how an RBP finds and binds to its targets. Finding sequence motifs is not trivial due to the shortness of the binding motif and the large number of input sequences that can include many false positives. Incorporating secondary structure preferences into motif models adds an extra layer of challenges due to the noisiness of RNA structure prediction and the need of a reliable model for sequence-structure motifs that is also easy to interpret. The RBP binding motifs can be derived either with methods developed for DNA-binding proteins, which consider only the RNA primary sequence, or with specifically designed tools that account for different levels of secondary structure information. The first motif finders designed for RBPs used RNA secondary structure as prior knowledge to restrict the search for sequence motifs to either single-stranded regions or to specific loop structures (Hiller *et al.*, 2006; Foat and Stormo, 2009; Li *et al.*, 2010). More recent tools use different strategies to model and predict both sequence and structure motifs. *RNAcontext* (Kazan *et al.*, 2010) detects the relative preferences of an RBP for multiple structural contexts. It uses a probabilistic framework to model separately the sequence and structure preferences, and was designed to work with RNAcompete binding-affinity data. Although it was also applied to CLIP datasets, its performance in this case is not established. *GraphProt* (Maticzka *et al.*, 2014) learns sequence and structure binding preferences of RBPs by modeling the binding sites as hypergraphs. It uses graph kernel-based support vector machines (SVM) to classify between bound and unbound regions. The predicted bound model is hard to interpret or visualize, and the tool outputs the top-scoring 1000 sequences and structures, that can be converted to PWMs or logos. *Zagros* (Bahrami-Samani *et al.*, 2015) is an extension of the MEME algorithm designed for CLIP data. It accounts for cross-link modification events and for secondary structure in the form of paired-unpaired probabilities. *Zagros* uses as input only the binding sites derived from experimental data and performs *de novo* motif discovery. These motif finders are designed specifically for a certain type of experimental data and only *Zagros* finds *de novo* sequence-structure motifs, while *RNAcontext* and *GraphProt* work in classification or regression settings. Furthermore, the structure predictions of all tools were not objectively evaluated.

In this article, we introduce **SSMART** (sequence-structure motif analysis tool for RNA-binding proteins), an RNA motif finder that extends cER-MIT (Georgiev *et al.*, 2010) – a sequence-based motif finder used primarily to determine DNA binding preferences from high throughput data such as CHIP-seq. Our tool identifies binding motifs by simultaneously modeling the primary sequence and the secondary structure of the RNA and searching for optimal sequence-structure motifs of flexible lengths. The sequence-structure motifs are represented as consensus strings over a degenerate alphabet, extending the IUPAC codes for nucleotides to also reflect secondary structure preferences. The secondary structure is obtained in a prior step, by sampling suboptimal structures around binding sites and identifying local dominant combinations of base pairs (Rogers and Heitsch, 2014). The motif candidates are evaluated with an objective function that integrates the individual RNA targets binding evidence into a combined score. The objective function is optimized with a greedy search strategy that starts with a set of 4-mers over the non-degenerate sequence-structure alphabet. Each of this “seed” motifs is then “evolved” iteratively until the motif score cannot be improved anymore. After all the motif seeds are evolved, **SSMART** applies a post-processing step in which the evolved motifs are clustered and corresponding PWMs are generated from high scoring candidates (Fig 1C).

**Figure 1:**
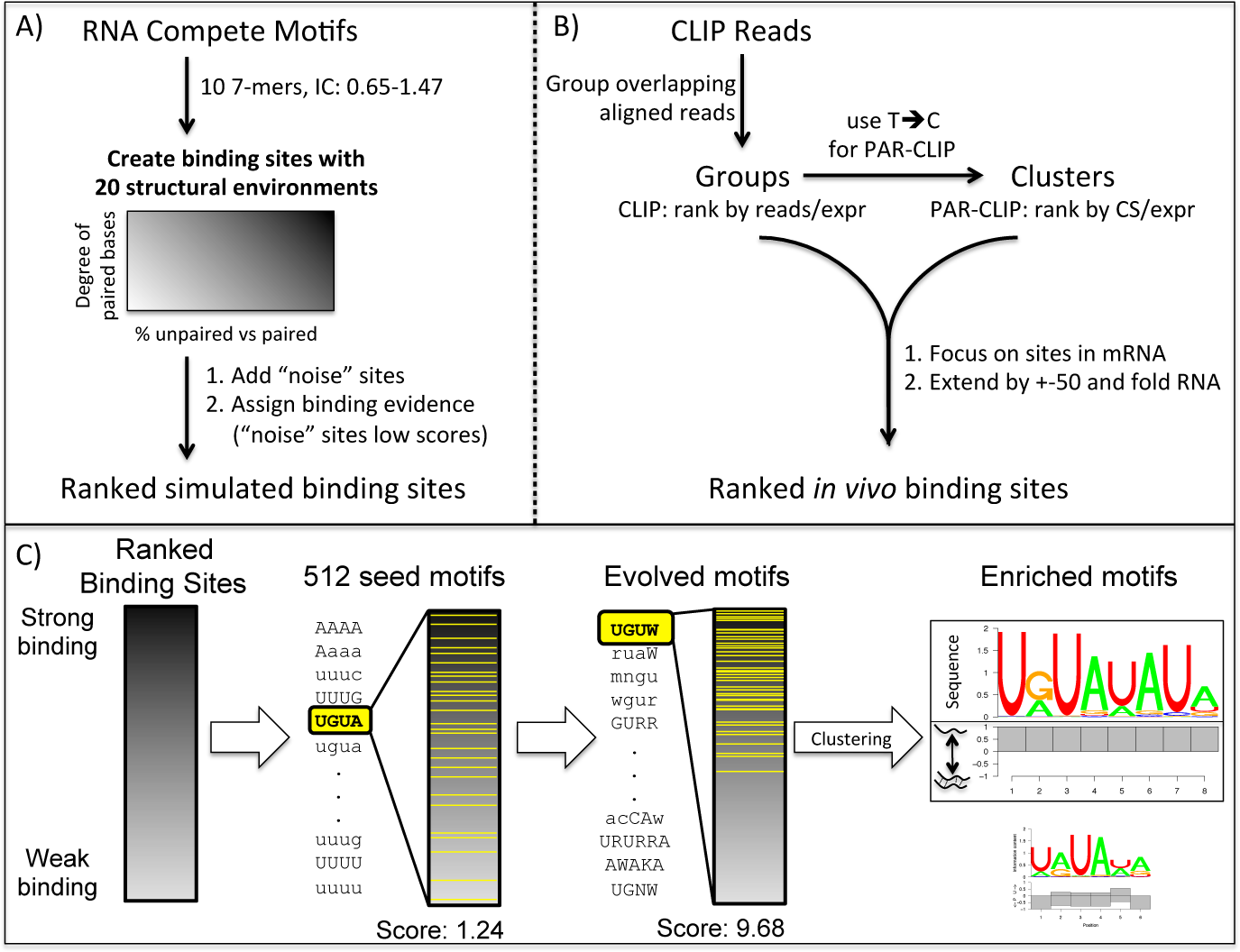
Overview of the motif finder workflow. A) Preparation of synthetic datasets. B) Pre-processing of CLIP reads. CS refers to T-to-C conversion specificity, and expr to gene expression. C) **SSMART** algorithm. A set of 4-mer seeds are independently evolved in order to optimize the score over the ranked list of binding sites. Then the motifs are clustered and the best sequence-structure motifs are reported. We represent the two components separately: the upper part corresponds to the sequence logo, while the lower part depicts the probability to be paired (below the line) and unpaired (above the line) for each base.

Evaluations on synthetic data showed that **SSMART** is able to recover RBP sequence and structure motifs implanted into 3‘UTR-like sequences, in various proportions of structured/unstructured binding sites. We successfully used **SSMART** on high-throughput *in vivo* and *in vitro* data, showing that we not only recover the known sequence motif, but also gain insight into the structural preferences of the RBP.

## 2 Methods

In this section we describe in detail the motif finding strategy implemented in **SSMART** as well as the employed evaluation procedures. We also explain the datasets that were used to test our tool, including how the synthetic ones were generated.

### 2.1 RNA secondary structure prediction

RNA molecules are flexible oligonucleotides that can adopt multiple stable structures. Their folding is influenced not only by their composition and the local environment, but also by other molecules, so it is difficult to obtain the exact secondary structure that an RNA has during an interaction with a protein *in vivo*. The available structure prediction tools are based either on free energy minimization (Zuker, 2003; Bernhart *et al.*, 2006), or on ensembles of secondary structures (Ding and Lawrence, 2003; Rogers and Heitsch, 2014). The more recent algorithms focus on local conformations and can take into account multiple suboptimal structures.

We considered two folding algorithms: RNAplfold (Bernhart *et al.*, 2006) and RNAprofiling (Rogers and Heitsch, 2014). RNAplfold is a tool from ViennaRNA package that predicts RNA single-strandedness using free energy minimization and locally stable secondary structures. It has two important parameters: the size of the window (*W*) and the maximum base pair span (*L*). RNAplfold associates the best structure to each sliding window over the stretch of the RNA of interest, and then outputs the average base pair probabilities. RNAprofiling is an ensemble-based method that balance abstraction and specificity by identifying local dominant combinations of base pairs. It uses a statistical sample of 1000 RNA secondary structures from the Boltzmann ensemble of possible RNA secondary structures associated with a given RNA sequence. The tool then focuses on the arrangement of helices at the substructure level and reports the most frequent double-stranded regions. Extensive testing revealed a strong length dependency for RNAplfold structures (bigger parameter values yielding more paired bases), while RNAprofiling results were stable (see Supplementary Section S1.1). We used RNAprofiling for all **SSMART** results reported here, but the user can compute secondary structures with any tool, if then the predicted structures are properly encoded into the input sequences.

### 2.2 The sequence-structure motif identification framework

In order to simultaneously model the primary sequence and the secondary structure of the RNA, **SSMART** represents the sequence-structure motifs as consensus strings over an extended degenerate alphabet. We use the regular IUPAC codes for nucleotides to denote bases in single-stranded positions, and their lower-case counterparts to denote bases in double-stranded context. Given a set of putative RBP binding sites with corresponding binding scores for each site, **SSMART** searches for optimal sequence-structure motifs of flexible lengths (Fig 1C). The framework has two essential components: an objective function that scores the binding strength of a given *k*-mer and a search procedure that explores the motif space for high-scoring *k*-mers. A **SSMART** motif of length *k* is a *k*-mer over the alphabet *A*_*complete*_ = {A, C, G, T, W, K, R, Y, S, M, N, a, c, g, t, w, k, r, y, s, m, n}. We define the motif space to be all *k*-mers with length between 4-10 over the *A*_*complete*_ alphabet, with a limited number of degenerate positions.

#### 2.2.1 Binding evidence

**SSMART** input consists in the set of *n* input sequences *s*_*i*_, *i* = 1, …, *n* (for example, CLIP peaks or RNAcompete oligos), described with the sequence-structure 8 letters alphabet *A*_*basic*_ = {A, C, G, T, a, c, g, t}, and their corresponding binding scores *y*_*i*_, *i* = 1, …, *n*. These scores depend on the type of experiment used to derive the binding specificities of the RBP in question. In the case of CLIP experiments, we used PARalizer peaks together with cell line-specific gene expression from RNA-seq data to define the following binding scores: normalized read counts 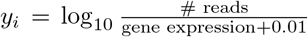 for CLIP-seq datasets; and normalized T-to-C conversion specificity

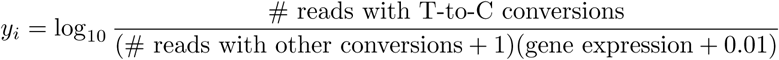

for PAR-CLIP datasets. For RNAcompete datasets we used the affinity scores (normalized signal intensities) to describe the binding preferences. We note that in the case of *in vivo* CLIP experiments the majority of sequences in the dataset correspond to binding events, while for *in vitro* RNAcompete data the majority of input sequences will be unbound. **SSMART** is able to handle both types of score distributions.

#### 2.2.2 The objective function

There are two available approaches for evaluating the binding strength of a given *k*-mer in our framework: a random set score and a linear regression score.

The random set approach (RS score) was described in Georgiev *et al.* (2010) and works well with a variety of scores that reflect the direct binding evidence, but is restricted by the assumption of independent contributions for the space of the input sequences. Given the sequences *s*_*i*_ and scores *y*_*i*_, a motif *m*_*j*_ partitions the sequence space into a positive set (containing *m*_*j*_) and a negative set (not containing *m*_*j*_). We search over the discrete space of possible motifs for the optimal motif *m*\* that yields high binding scores in the positive set and low binding scores in the negative set. Given a motif *m*_*j*_, we denote the number of sequences with motif occurrences with 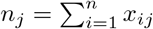, where the binary variable *x*_*ij*_ indicates a match of *m*_*j*_ in sequence *s*_*i*_. We consider the enrichment score 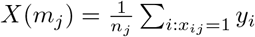 as a random variable whose randomness comes through the set of sequences containing *m*_*j*_ (*x*_*ij*_ = 1), and not through the scores *y*_*i*_, and we define the random set scoring function to be its z-score:

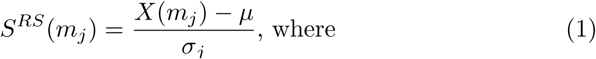

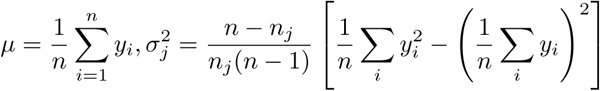

This is a zero mean, unit variance test statistic on the null hypothesis that sequences containing *m*_*j*_ are not enriched for the motif *m*_*j*_. The optimization problem is:

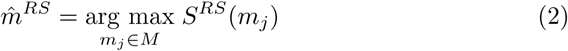

where *M* is the set of putative motifs {*m*_1_, …, *m*_*p*_, and 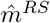 is the best guess at the optimal binding motif *m*\*.

We can extend the scoring function to any rule *R*(*s*_*i*_) that partitions the sequence space into two sets as follows: We denote by *x*_*iR*_ the truthfulness of rule *R* in sequence *s*_*i*_. Then 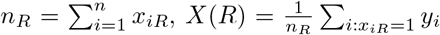, and the induced score is *S*^*RS*^(*R*) = (*X*(*R*) − *µ*)/*σ*_*R*_. The previous score is a particular case, with the rule *R*(*s*_*i*_) = (*m*_*j*_ ⊂ *s*_*i*_).

The linear regression approach (LR score) was introduced in Corcoran *et al.* (2011) and is more computationally demanding but can account for some, potentially relevant, confounder information, like di-nucleotide frequencies or sequence length (see Supplementary Section S1.2).

#### 2.2.3 The search strategy

We need to search the sequence-structure motif that optimizes one of the objective functions defined before (Eq. 1 or Supplementary Eq. 8). An exhaustive search over the space of all potential motifs is not computationally feasible, thus we employ a custom greedy search strategy that considers a large set of seed motifs that are independently updated. These seed points are motifs of length 4 with the same structure and all possible sequence composition (512 4-mers over the alphabets {*A, C, G, T*} and {*a, c, g, t*}). We note that we obtain consistently similar results with this (reduced) set as with the whole set of 4096 possible 4-mers over the *A*_*basic*_ alphabet.

Given a motif *m*, a set of candidate motifs is constructed by applying small variations to *m*: in length, sequence, or structure. The *k*-mer *m* is extended to 16 new (*k* + 1)-mers, by independently adding one letter from *A*_*basic*_ at one of its end. If *k >* 4, the length of the motif is reduced and 2 new (*k* − 1)-mers are considered. Then a large set of new *k*-mers are obtained by changing one letter at a time in terms of structural change or increasing/decreasing sequence degeneracy (see Supplementary Section S1.3).

Each seed motif *m*_*i*_ starts an independent search for the best motif. At one iteration, all update rules are applied and each new motif candidate is scored. The motif candidate with the highest motif score is used in the next iteration. The procedure is repeated until the motif score cannot be improved, in which case the last motif is reported. The result of the search is a set of evolved motifs 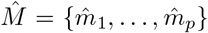 and their corresponding scores 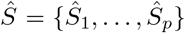. For each motif 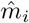, its occurrences in the top 50% input sequences are used to derive a PWM.

#### 2.2.4 Post-processing procedure

The complete set of evolved motifs will have many similar motifs that vary by a few letters or have different lengths and/or overlap (see Supplementary Section S1.4). **SSMART** applies a post-processing procedure in order to cluster multiple evolved motifs together and to rank these merged motifs. We use the metric introduced by Harbison *et al.* (2004) to define a similarity measure as follows. For two motifs *a*, *b* of equal length *w*, the Harbison distance is 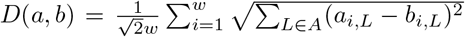, where *A* is the alphabet and *a*_*i,L*_, *b*_*i,L*_ are the probabilities of observing base *L* at position *i* of motifs *a* and *b*, respectively. As in Georgiev *et al.* (2010), we define the following similarity score:

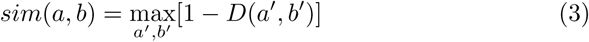

where *a*′, *b*′ correspond to all possible overlaps between motifs *a*, *b* induced by shifts such that the minimum overlap length is 3.

Given the set of redundant output motifs 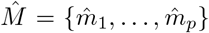, we obtain a set of ordered motif clusters {*C*_*i*_} with the following clustering procedure:

1. Initialize the cluster count: *q* = 1;
2. Find the top motif in 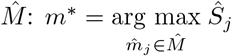
3. Select all motifs 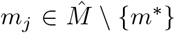 with *sim*(*m*\*, *m*_*j*_) *≥* 0.75 and compute scores for the union rule *R*_*|*j*_(*s*_*i*_) = (*m*\* ⊂ *s*_*i*_)|(*m*_*j*_ ⊂ *s*_*i*_) and the intersection rule *R*_*&*j*_(*s*_*i*_) = (*m*\* ⊂ *s*_*i*_)&(*m*_*j*_ ⊂ *s*_*i*_);
4. Add *m*\* and all similar motifs *m*_*j*_ that have *S*(*R*_*|*j*_) *≥* 0.95 ⋅ *S*(*m*\*) or *S*(*R*_*&*j*_) *≥* 0.95 ⋅ *S*(*m*\*) to *C*_*q*_;
5. Remove the set *C*_*q*_ from 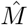;
6. Update cluster count: *q* = *q* + 1;
7. Repeat steps 2 to 6 until 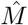 is empty.

For each motif cluster *C*_*i*_ we compute an aggregate PWM by averaging the PWMs of each cluster member weighted by its motif score.

### 2.3 Synthetic datasets

We generated synthetic datasets that contain specific implanted motifs in various proportions of structured/unstructured binding sites (Fig 1A). First, we selected 10 PWMs derived from RNAcompete experiments from the RBP compendium (Ray *et al.*, 2013). They all have length 7, but have different nucleotide compositions and their average information content varies between 0.65 and 1.47 (see Supplementary Table S1). We then used a 2nd order Markov chain to generate a large set of 500.000 random 3‘UTR sequences with lengths following the empirical distribution observed in PAR-CLIP peaks. In order to obtain different structural environments, each PWM was implanted into all synthetic 3‘UTR sequences in random locations, and then the secondary structure was predicted. Based on the number of predicted unpaired bases of the implanted motifs, we considered 20 different structural combinations, A-T (see Supplementary Fig S4). Structures A-K represent linear combinations of purely single-stranded and double-stranded binding sites, from A with 100% unpaired motifs, to K with 100% paired motifs (with 10% increments). Structures L-Q represent various degrees of double-strandedness in the binding, from set L with all sequences having 1 paired base, to structure Q with 6 paired bases in the implanted motif. The last 3 structural environments (R-T) denote variable structures: R has 30% unpaired motifs and 10% of each set with 1 to 7 paired bases, S has equal numbers (12.5%) of motifs with 0 to 7 paired bases, and T has 35% paired motifs, 35% unpaired motifs and 5% of each of the rest. For each implanted motif and each structural environment, we randomly selected 10 datasets of 2000 sequences each, generating a total of 2.000 synthetic datasets. We then added some noise to this data as follows: we generated a single set of 10000 3‘UTR sequences with the same 2nd order Markov chain, and then we predicted the corresponding secondary structure. In each synthetic dataset, we inserted 500 sequences selected at random from this “noise” set. The resulted datasets represented the “core” data for our comparison.

Since **SSMART** requires as input a binding score for each sequence, we sampled 2500 such scores (conversion specificity) from 25 PAR-CLIP datasets. Then we randomly associated these values to sequences in the generated datasets, making sure that the 500 “noise” sequences will be triangularly distributed among the positive sequences (less at the top, more at the bottom). For *Graph-Prot*, we generated a “negative” set of 2500 sequences with 3‘UTR composition.

### 2.4 Experimental datasets

We applied **SSMART** on high-throughput *in vivo* and *in vitro* data from CLIP and RNAcompete experiments, respectively. We selected and analyzed 10 different proteins: ELAVL1, FMR1, FUS, IGF2BP2, IGF2BP3, LIN28A, QKI, SRSF1, SRSF7 and SRSF9 that have both types of data available. We added two more proteins that had only CLIP data (see Supplementary Table S2). We retrieved the selected RNAcompete datasets from Ray *et al.* (2013). We downloaded the CLIP datasets from Gene Expression Omnibus (Edgar *et al.*, 2002) and processed the reads as described in Mukherjee *et al.* (2014) (see Fig 1B). Briefly, the reads from each library were pre-processed and aligned to the corresponding genome (hg19, mm10) and then interaction sites were defined with PARalyzer (Corcoran *et al.*, 2011). For all PAR-CLIP datasets we considered all PARalyzer clusters that corresponded to mRNAs and to each we associated the T-to-C conversion specificity normalized by gene expression, while in the case of CLIP-seq we used the groups with the normalized read counts. In order to obtain more realistic structure predictions, we extended each binding site by maximum 50 bp on each side using either the genome (for the intronic regions) or the transcriptome (for the rest). For each cell line we derived transcript abundance from RNAseq data, and then for each considered site we retrieved the flanks up to 50 bp from the most abundant transcript that contained it.

### 2.5 Evaluation on synthetic datasets

In order to evaluate the motif finders performance we converted all sequence and structure predictions to a uniform encoding. For sequence motifs we used PWMs, converting *RNAcontext* energy matrices and *GraphProt* list of top 1000 sequences to probability matrices. For **SSMART** we collapsed the predicted PWM over the 8 letter extended alphabet to a 4 letter alphabet. In the case of structures, *Zagros* and **SSMART** use two structural contexts per nucleotide (paired and unpaired) while *RNAcontext* and *GraphProt* use larger but distinct sets of structures (e.g. stem, hairpin loop, internal loop, etc.), thus we converted the predicted structures of all tools to a vector of paired probabilities.

We evaluated the motif finders performance on synthetic datasets by computing the recovery rates for sequence and structure motifs separately. For all tools we considered one motif, taking the top one when more motifs are reported. We compared the recovered motifs with the implanted motifs by computing similarity scores based on the Harbison metric (used also in the post-processing step). We use Eq. (3) to define the similarity score between two sequence motifs *a* and *b*, with the mention that *a*′, *b*′ correspond to all possible overlaps between motifs *a*, *b* induced by shifts such that the minimum overlap length is max(*w* − 1,4), where *w* is the smaller motif length. We also define the similarity score for the corresponding structures *s*,(*a*) and *s*(*b*) to be: *sim*(*s*(*a*), *s*(*b*)) = 1*−D*(*s*(*a*′), (*b*′)), where 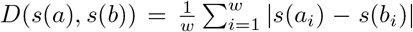. For each tool, we computed its threshold for “recovered” and “not recovered” motifs by comparing each of the 2000 predicted motifs with all 200 implanted motifs. Then the optimal cutoffs for sequence motifs and for structure motifs are determined independently by optimizing the p-values obtained with G-tests of independence (see Supplementary Table S5 and Supplementary Figure S5) (Sokal and Rohlf, 2012).

### 2.6 Evaluation on CLIP datasets

We compared **SSMART**, *GraphProt* and *Zagros* on 6 selected PAR-CLIP libraries corresponding to 2 proteins: ELAVL1 (HuR) and PUM2 (see Supplementary Table S4). For each tool and library we retrieved the predicted sequence motif in the form of a PWM, and the sequence-structure motif as a PWM over the 8 letter extended alphabet. Then we tested how well a motif predicted on a particular library correlates with the binding scores associated with each of the 6 considered libraries. We used the Kendall tau correlation coefficient between the ranked list of binding scores and the corresponding log-likelihood scores of a given PWM. We note that the tau coefficient has values in [−1, 1], a value close to 1 indicating strong agreement, while a value close to −1 indicating strong disagreement. For each tool and protein, we then applied a two-sample Kolmogorov-Smirnov test, comparing the tau correlations on datasets for the same RBP versus those on the other protein.

## 3 Results

**SSMART** performs *de novo* motif discovery on high-throughput RNA-binding protein data, predicting sequence and structure binding motifs of RBPs. We generated synthetic datasets with certain motifs in different structural context in order to evaluate its performance and to compare the prediction of sequence and structure motifs with three other RBP sequence-structure motif finders: *RNAcontext*, *GraphProt* and *Zagros* (Kazan *et al.*, 2010; Maticzka *et al.*, 2014; Bahrami-Samani *et al.*, 2015). We also compared **SSMART** with *GraphProt* and *Zagros* in a cross-validation setting across replicate *in vivo* CLIP libraries. We then used **SSMART** to examine a range of publicly available biological datasets and to compare binding specificities derived from *in vivo* and *in vitro* experiments. Afterwards we analyzed the structural binding specificity for a selection of CLIP datasets. In this section we present the results of our analyses.

### 3.1 Recovering sequence and structure motifs from synthetic datasets

Evaluation of *de novo* motif predictions from experimentally-derived datasets is challenging due to lack of a known ground truth and noise. Therefore, we generated a large set of datasets that mimic PAR-CLIP binding sites (clusters) in which we inserted 10 different motifs derived from RNAcompete experiments. For each motif we considered 20 different structural combinations (A-T) and we measured not only how well each tool recovers it, but also how well the initial structure is predicted.

The recovery rates for sequence and structure motifs on all 2000 datasets are presented in Table 1. Our tool outperforms all other motif finders in recovering the structure and is outperformed by *GraphProt* and *Zagros* in the sequence recovery, but has the best results for the combination of both sequence and structure. Although *Zagros* recovers 100% of the sequence motifs, its structural predictions are on the same scale as **SSMART**-seq, which considers all motifs to be single-stranded.

**Table 1:** Global recovery rates for sequence and structure motifs on synthetic datasets. The values reported for SSMART-seq correspond to a version of **SSMART** that uses only sequence information.

| Tools | SSMART | SSMART-seq | RNAcontext | GraphProt | Zagros |
| --- | --- | --- | --- | --- | --- |
| Sequence | 91.75 | 92.05 | 58.14 | 94.84 | 100 |
| Structure | 88.65 | 14.75 | 50.03 | 73.25 | 15.25 |

The sequence predictions performance is consistent for all tools across different structural environments (Fig 2A). We note that the motif appears to have some influence on the prediction performance for some motif finders, for example **SSMART** recovering the sequence motif with the lowest information content in just 46.5% datasets or *GraphProt* recovering the ACAACRR motif in 58% cases. In the case of structure predictions, all the tools exhibit variability across structural environments. **SSMART** and *RNAcontext* perform better on sets with more defined structures, while *GraphProt* recovers the mixed structures. This difference is explained by the way each tool models and reports the motifs. **SSMART** can capture mixed structures by reporting 2 or more separate motifs, but in this settings we consider only the top reported motif. Even so, **SSMART** recovers perfectly the structure if it has 80-100% of the binding sites in either unpaired or paired states (sets A-C and I-K) or if it has 6 of the 7 bases in the same structural context (sets L and Q). If just 5 bases have the same structure, the recovery rates are 98% and 94% for unpaired and paired RNA, respectively. In summary, **SSMART** recovers more than 90% of structural motifs for 15 (out of 20) structural environments, while *GraphProt* for only 7 types of structures.

**Figure 2:**
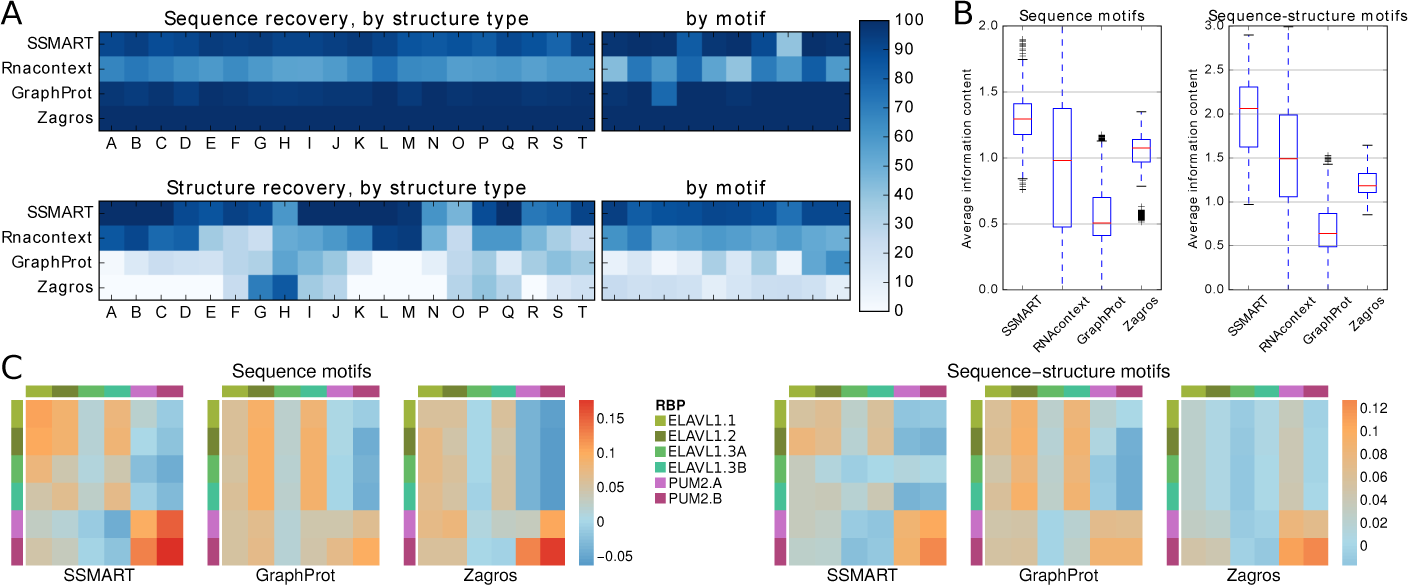
Comparison with other tools on synthetic and biological datasets. (A) Recovery rates on synthetic data for sequence motifs (top) and for structure motifs (bottom). The colors represent the percentage of recovered motifs from the datasets grouped by structure type or by implanted motif. (B) Average information content for predicted motifs on synthetic data, either in their sequence component (left) or for combined sequence-structure motifs (right). (C) Kendall tau correlation coefficients between the motifs predicted on one specific CLIP dataset versus the binding scores of a list of CLIP libraries. The correlations are depicted with the same color scale for the sequence motifs (left) and the combined sequence-structure motifs (right). The rows correspond to the training sets, and the columns to the test sets.

Next, to assess the specificity of identified motifs relative to background, we computed the average information content of the predicted motifs (Fig 2B). We considered two variants of motif information content: for sequence motifs we used average information content over the 4 letter sequence alphabet *A* = { A,C,G,T}; for sequence-structure motifs we derived average information content using the 8 letter sequence-structure alphabet *A*_*basic*_ = {A, C, G, T, a, c, g, t}. In the case of sequence motifs, the median information content was 1.29 for **SSMART**, 0.98 for *RNAcontext*, 0.5 for *GraphProt* and 1.07 for *Zagros*. *RNAcontext* sequence motifs cover the whole range of possible information content, while the values for *Zagros* have the smallest variance. The low value for *GraphProt* is explained in part by the length of 12 bases reported for all motifs. **SSMART** produces the most expressive sequence-structure motifs, with a median information content of 2.06. The values for *RNAcontext*, *Zagros* and *GraphProt* are 1.49, 1.18 and 0.64 respectively. Taken together these results demonstrate that **SSMART** provides the best all-around performance on the simulated data.

### 3.2 Testing motif predictions on CLIP datasets

Next, we used published CLIP datasets to evaluate the motif predictions of **SSMART**, *GraphProt* and *Zagros* in a train/test setting, by correlating each learned motif with the binding scores of all considered libraries. A meaningful motif will exhibit positive correlation for libraries of the same protein and negative or smaller correlation coefficients for inter-protein tests. The Kendall tau correlation coefficients obtained for 4 ELAVL1 (HuR) and 2 PUM2 PARCLIP libraries are presented in Fig 2C. The datasets denoted with A and B are replicates, while the ones denoted 1, 2, and 3 are from independent experiments. ELAVL1.3 CLIP was performed in HeLa cell line, while the rest in HEK293 cells. For all tools, the sequence motifs trained on the ELAVL1 datasets (first 4 rows) perform as expected, with negative or lower values obtained when tested on PUM2 binding sites, the corresponding p-values being bellow 0.0001 (see Supplementary Table S5). The only outlier is the ELAVL1.3A dataset, on which all tested motifs obtain lower correlations. However, **SSMART** is the only motif finder that shows the same trend not only in the ELAVL1.3A column, but also in the ELAVL1.3A row. On the other hand, the sequence motifs trained on the PUM2 datasets (last 2 rows) are protein-specific only in the case of **SSMART**, with a p-value of 0.002, while for *GraphProt* and *Zagros* the predicted motifs correlate similarly with PUM2 and ELAVL1 binding sites (p-values of 0.335 and 0.061, respectively).

### 3.3 Identification of motifs from *in vivo* and *in vitro* datasets

We applied **SSMART** to 36 CLIP and 21 RNAcompete libraries corresponding to 12 proteins with both *in vivo* and *in vitro* experiments. For the full list of results see Supplementary Sections S3 and S4. All RNAcompete data is from human and was downloaded from the compendium of RNA-binding motifs (Ray *et al.*, 2013). The *in vivo* data corresponds to 31 PAR-CLIP and 5 CLIP-seq experiments conducted in human HEK293, HeLa, and H9 hESC cells; three of the CLIP-seq datasets were performed in A3 lymphocytes or mESC cells.

First, we compared RBPs for which both *in vivo* and *in vitro* experimental data was available (Fig 3). For each RBP we present the top sequence-structure motifs for CLIP and RNAcompete datasets, as well as the motifs reported by the authors of the respective experiments. We note that in the case of RNA-compete experiments, **SSMART** derives the binding motif from all ~240,000 probes with corresponding affinities, while Ray *et al.* (2013) derived enrichment scores for all possible 7-mers and then defined motifs from the top 10 7-mers. Nevertheless, we find strong agreement between our top predictions and the reported RNAcompete motifs for almost all RBPs examined. None of the *in vitro* results indicated any structural features, which may be due to the length and/or selection of RNA oligos in the RNAcompete assay.

**Figure 3:**
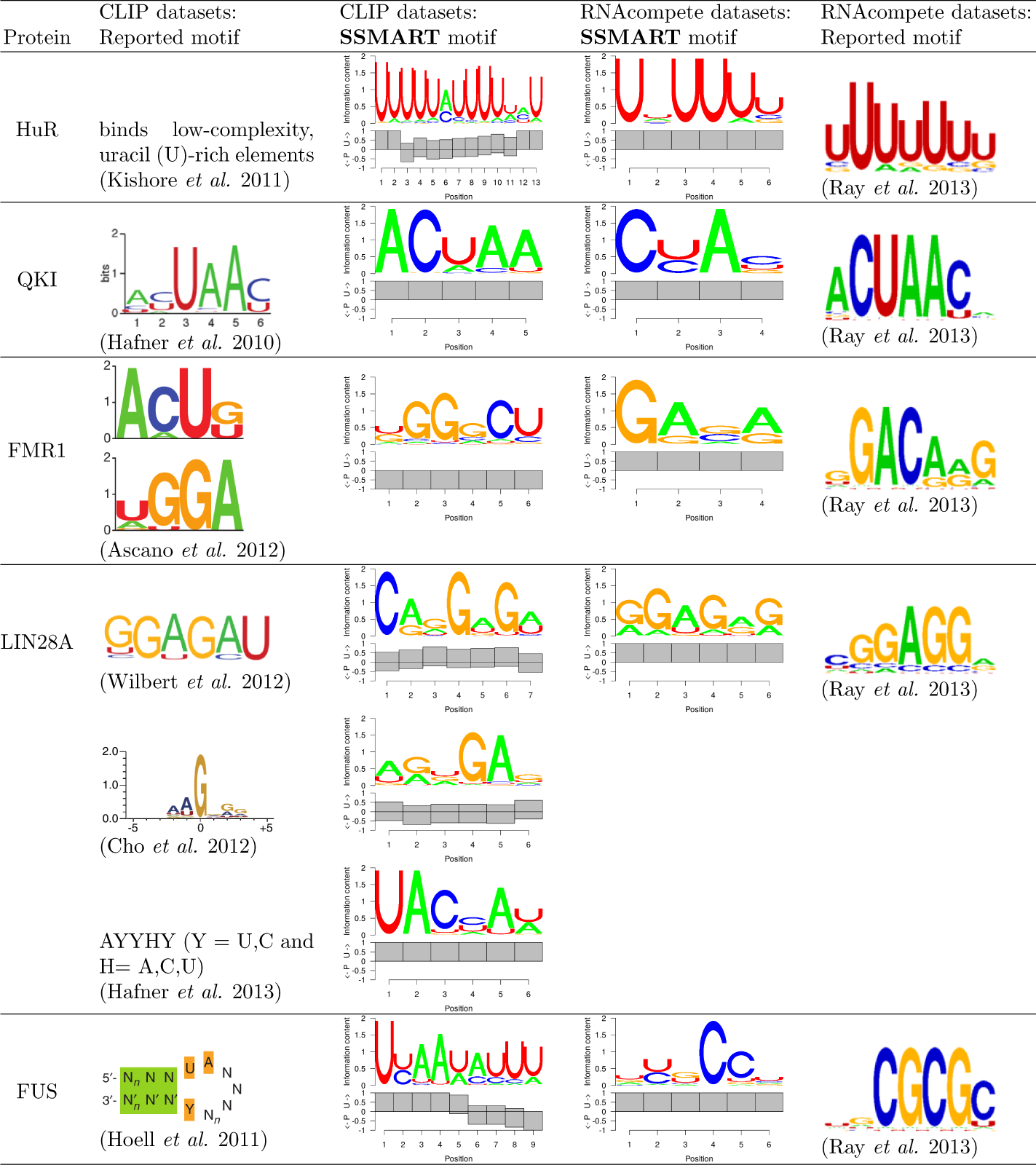
**SSMART** results on biological data: *in vivo* vs. *in vitro* comparison.

RBPs exhibited varying degrees of concordance between A) predictions and results for *in vitro* data, B) predictions and reported results for *in vivo* data, and C) predictions and results for *in vivo* and *in vitro* data. For both ELAVL1 (HuR) and QKI, we observed full agreement (i.e. *in vivo* and *in vitro* predictions and results identified the same motif). For ELAVL1, the U-rich sequence motif recovered by **SSMART** from *in vivo* data was associated with mostly single-stranded structural context As expected (Feracci *et al.*, 2016), the motifs reported for QKI were associated with single-stranded structural context.

Fragile X-mental retardation 1 (FMR1) is a RNA-binding protein that has multiple distinct RNA-binding domains. PAR-CLIP experiments reported two short binding motifs, ACUK and WGGA, that interact with the KH and RGG domains, respectively (Ascano *et al.*, 2012). Our top predicted motif is similar to WGGA and associated with paired RNA, which may reflect previously reported binding to G-quadreplex structures (Brown *et al.*, 2001). A CU di-nucleotide, which is present in the secondary ACUK motif, was present in the top scoring motif. The *in vitro* predicted and reported results did not identify the secondary motif bound by the KH domain, nor did it indicate a structural context for the WGGA.

The *in vitro* prediction for lin-28 homolog A (LIN28A) was consistent with the reported motif from RNAcompete. Similarly, our *in vivo* predictions from LIN28 CLIP-seq data from Yeo and Kim labs identified a GA-rich motif consistent with what was reported (Cho *et al.*, 2012; Wilbert *et al.*, 2012). Both the predicted and reported results from LIN28A PAR-CLIP were not consistent with the *in vitro* results or the *in vivo* CLIP results. Also, for some analyzed proteins the *in vitro* predictions were in agreement with the reported compendium motif, but the predicted motifs for *in vivo* datasets showed different binding specificities (data not shown). The basis for the inconsistency is unclear and could be due to technical differences between CLIP and PAR-CLIP, ranking and normalization of called peaks, or the cell lines the experiments were performed in.

FUS was a clear case in which there was concordance between our predictions and reported results both *in vivo* and *in vitro*, however the reported *in vivo* and *in vitro* specificities differ. The *in vitro* results indicate a CG-rich motif, while the *in vivo* results suggest a UA-rich sequence, which has been reported to have structural context (Hoell *et al.*, 2011), which we will describe in more detail below. This may represent an example in which the *in vitro* results may not accurately reflect *in vivo* binding.

### 3.4 Examining the structural context of binding specificity

Due to the lack of structural insights from the RNAcompete results, we focused on *in vivo* predictions exhibiting markedly different structural context for a subset of RBPs (Fig 4). A stem-loop structure with some sequence preference was previously reported for FUS (Hoell *et al.*, 2011). The top ranked motif predicted for the FUS PAR-CLIP data was consistent with the reported binding preference in both sequence and structure. From 5’ to 3’ the predicted motif is decreasingly single-stranded particularly with an apparent transition from unpaired to paired at position 6, presumably representing the loop, with the reported UA at the begining of the loop (positions 4-5).

**Figure 4:**
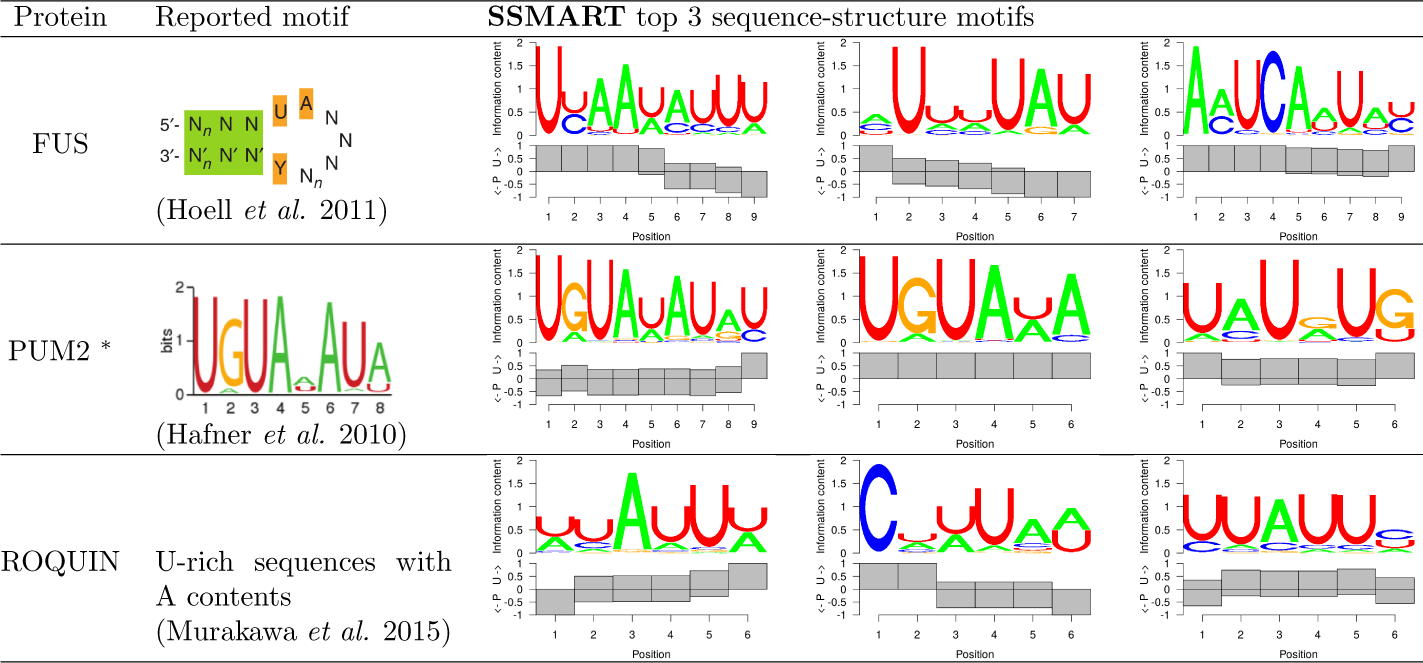
**SSMART** top 3 sequence-structure motifs for three selected CLIP datasets. * marks predictions obtained with the random set scoring.

Examination of Pum2 PAR-CLIP data revealed the well-established UGUA-HAUA binding motif (Hafner *et al.*, 2010). As expected, we found this motif in a single-stranded context (Lu and Hall, 2011). Interestingly, we also identified the motif in a paired context, which may represent sites in which modulation of secondary structural switch influences Pum and miRNA-mediated regulation (Kedde *et al.*, 2010).

Examination of Roquin (RC3H1) binding specificity using PAR-CLIP did not reveal a specific sequence motif, however they proposed existence of a stemloop with an AU-rich loop region (Murakawa *et al.*, 2015). The top predicted motif match is consistent with the reported sequence description. This prediction could represent, predominantly, the loop portion of the proposed stem-loop. The results described for these three RBPs highlight the manner in which incorporation of secondary structure can enhance the interpretation of RBP-binding specificity, particularly for *in vivo* experimental data.

## 4 Discussion

We developed **SSMART**, a *de novo* motif finder that identifies sequence-structure binding motifs from large sets of RNA sequences derived from genome-wide *in vivo* or *in vitro* experiments such as CLIP or RNAcompete. Our tool simultaneously models the primary sequence and the structural properties of the RNA target sites and produces easy to interpret sequence-structure binding motifs. **SSMART** searches for optimal sequence-structure motifs of flexible length in putative RBP binding sites ranked by their experimentally-derived binding evidence. While *Zagros* and *GraphProt* were designed for CLIP data and *RNA-context* is best suited for RNAcompete data, our approach is more general and can successfully handle different types of input. Moreover, **SSMART** learns all motif characteristics from the data, including the motif length, and does not require parameter optimization. Like *Zagros*, **SSMART** accounts only for double-stranded and single-stranded preferences at each individual position of the motif. In contrast, *RNAcontext* and *GraphProt* distinguish between five different structural contexts, but they output aggregate structural motifs. Our tool is able to identify different sequence motifs with the corresponding per base structural preference for the same protein.

Although it was reassuring that **SSMART** performed well on simulated and *in vitro* data, ranking binding evidence is straightforward in these scenarios, unlike for *in vivo* binding sites. RNA expression levels clearly impact the read-evidence and there are other factors, such as cross-linking or RNase choice, which may need to be incorporated to properly rank *in vivo* binding sites. Therefore input or background binding libraries may be more useful, particularly for intronic binding sites for which RNA expression estimates could be problematic. Appropriate normalization and ranking of *in vivo* (i)CLIP or PAR-CLIP data remains an ongoing challenge in the field.

We identified cases in which *in vitro* and *in vivo* results were discordant. Biases in both *in vitro* and *in vivo* assays may explain these differences. However, these differences could also be due to factors influencing *in vivo* binding that cannot be recapitulated *in vitro*. Biologically relevant explanations include RNA structural constraints, multiprotein RNA-binding complexes, as well as biophysical features of RNP granules in which these interactions occur. Our results indicate that **SSMART** should assist investigators in accounting for RNA-structural constraints. Importantly, **SSMART** is general enough to be utilized as the determination of RNA-structure progress both experimentally and computationally.

In conclusion, we propose an efficient algorithm to identify the most probable sequence-structure motif, or combination of motifs, given a large set of RNA sequences. Our method can contribute to the systematic understanding of RBP-RNA binding specificity as more genome-wide experiments that determine RBP binding are performed.

## Acknowledgements

The authors would like to thank Stoyan Georgiev for his assistance in understanding the source code of cERMIT. Also, we would like to thank Dan Munteanu for his help in setting up some of the tools used.

