## Supplementary Materials for "SSMART: Sequence-structure motif identification for RNA-binding proteins"

### S1 Materials and Methods

#### S1.1 RNA secondary structure prediction

We considered two folding algorithms: RNAplfold (Bernhart *et al.*, 2006) and RNAprofiling (Rogers and Heitsch, 2014), and we compared their results on binding sites for two proteins with different structural preferences: PUM2, an RBP that prefers ss-RNA; and Staufen1, known to bind ds-RNA. We predicted RNA secondary structures for PUM2 and Staufen1 binding sites derived from *in vivo* PAR-CLIP and RIPiT data, respectively (Hafner *et al.*, 2010; Ricci *et al.*, 2014). We extended the core regions with maximum 25, 50, 75 or 150 nucleotides on each side, using either genomic coordinates in the case of intronic sequences, or the highest expressed isoform that contains the peak for the exonic ones. First, we compared RNAplfold predictions in multiple parameter settings, and then we compared the RNA secondary structure predictions of RNAplfold vs. RNAprofiling.

##### S1.1.1 RNAplfold predictions depend on parameters

RNAplfold is a tool from ViennaRNA package that predicts RNA single-strandedness using free energy minimization and locally stable secondary structures. It associates the best structure to each sliding window over the stretch of RNA of interest, and then outputs the average base pair probabilities. RNAplfold has two important parameters: the size of the window ($W$) and the maximum base pair span ($L$). Multiple applications of RNAplfold use the values $(W, L) = (80, 40)$ (see (Li *et al.*, 2010; Kazan *et al.*, 2010; Marin and Vanicek, 2011; Lekprasert *et al.*, 2011)), while (Lange *et al.*, 2012) recommend that $W = L + 50$ in order to have each base present in at least 51 windows.

We selected a wide range of parameters: $L \in [30, 150]$ and $W \in [L, L + 100]$, resulting in 143 $(W, L)$ pairs, and we applied RNAplfold on 3974 PUM2 and 4666 Staufen1 peaks. In the folding step, we extended the peaks with maximum 150 bp on each side, after which we discarded the flanking regions and we analyzed only the initial binding sites with the associated RNA structures. Fig. S1 shows the percent of bases that were predicted to correspond to ds-RNA (bases that have the predicted unpaired probability $< 0.5$). RNAplfold predicts on average 13% more paired bases for Staufen1 binding sites than for PUM2 binding sites, a result consistent with the reported binding preferences for the two RBPs. We note that in both sets the results show a clear trend of more base pairs as the parameter values increase. This is also in agreement with the reported behaviour of multiple structure prediction tools, namely that the prediction accuracy for individual base-pairs decreases with respect to span length and/or window length (Doshi *et al.*, 2004; Kiryu *et al.*, 2011;

Lange *et al.*, 2012). Given this correlation between the parameter size and the percent of paired bases for both sets, it is unclear what the "optimal" parameter values might be. Looking at just two widely used parameter values, $(W, L) \in \{(80, 40), (150, 100)\}$, we observe an unsettling difference of 11.1% for PUM2 and 9.3% for Staufen1 in the number of paired RNA bases.

###### S1.1.2 Comparative analysis

Given the performance of RNAplfold with different parameter settings, we looked for another tool for RNA structure prediction, and we selected RNAprofiling, which has a different approach to RNA folding. RNAprofiling is an ensemble-based method that balance abstraction and specificity by identifying local dominant combinations of base pairs. It uses a statistical sample of 1000 RNA secondary structures from the Boltzmann ensemble of possible RNA secondary structures associated with a given RNA sequence. The tool then focuses on the arrangement of helices at the substructure level and reports the most frequent double-stranded regions. We consider a base to be paired only if it is contained in a helix that is present in more than half of the ensemble of secondary structures. RNAprofiling has no parameters, the results being influenced only by the length of the input sequence. We tested it with three different sizes for the flanking regions of the RBP binding sites: 25, 50 and 75 and we compared the results with RNAplfold predictions for $(W, L) \in \{(80, 40), (150, 100), (200, 150)\}$.

We used the Paired/Unpaired predictions to define the following similarity metric between two foldings $a$, $b$ of the same sequence of length $w$:

$$Sim(a, b) = \frac{1}{w} \sum_{i=1}^{w} s_i, \text{ with } s_i = \begin{cases} 1, & \text{if } a_i \neq b_i \\ 0, & \text{if } a_i = b_i \end{cases} \tag{1}$$

where $a = \{a_1, a_2, \ldots, a_w\}$ and $b = \{b_1, b_2, \ldots, b_w\}$. A similarity score of 0 denotes identical structures, while a score of 0.3 means that the two predicted structures disagree in 30% of their positions. We compared the structures predicted by the two tools, each in three settings, for PUM2 and Staufen1 binding sites. The average similarity scores are depicted in Fig S2A. We note that the similarity between structure predictions for PUM2 (above the main diagonal) and Staufen1 (below the diagonal) has the same trend, with smaller values for comparisons between the same tool or between parameter settings with simmilar window lengths. The results that disagree the most (around 30%) correspond to the following pairs:

- RNAprofiling with shortest sequences (25 bp padding) & RNAplfold with the longest parameters ($W = 200, L = 150$), and
- RNAprofiling with longest sequences (75 bp padding) & RNAplfold with the shortest parameters ($W = 80, L = 40$).

The structural predictions that agree the most acros different tools (23-24%) correspond to simmilar folding sequences:

- short sequences: RNAprofiling with 25 bp padding & RNAplfold with $W = 80, L = 40$, and
- mid-range sequence: RNAprofiling with 50 bp padding & RNAplfold with $W = 150, L = 100$.

We also analyzed the percent of bases that were predicted to correspond to ds-RNA (Fig S2B). Staufen1 binding sites have an average 12% more paired bases than PUM2 sites, a trend consistent with the binding preferences shown by these RBPs. However, the secondary structure predicted by RNAplfold is correlated with the parameters used, longer values yielding more paired bases, while RNAprofiling predictions are quite stable with respect to the length of flanks.

We used RNAprofiling for all **SSMART** results reported here, but the user can compute secondary structures with any tool, if then the predicted structures are properly encoded into the input sequences. In order to predict secondary structures relevant to the experimental data, we apply the folding algorithm either on the RNA oligos in the case of RNAcompete data, or on extended peaks in the case of CLIP datasets (±50 bp). The extension is performed using either genomic coordinates in the case of intronic sequences, or the highest expressed isoform (from RNA-seq data) that contains the peak for the exonic ones. The extended RNA sequences are used only in the structure prediction step, after which only the core sequences are associated with the secondary structures in **SSMART** input files.

#### S1.2 The objective function in regression setting

The random set scoring function used by **SSMART** is defined by:

$$S^{RS}(m_j) = \frac{\frac{1}{n_j}\sum_{i:x_{ij}=1} y_i - \mu}{\sigma_j} \tag{2}$$

$$\mu = \frac{1}{n}\sum_{i=1}^{n} y_i, \sigma_j^2 = \frac{n-n_j}{n_j(n-1)}\left[\frac{1}{n}\sum_i y_i^2 - \left(\frac{1}{n}\sum_i y_i\right)^2\right]$$

The associated optimization problem is:

$$\hat{m}^{RS} = \arg\max_{m_j \in M} S^{RS}(m_j) \tag{3}$$

where $M$ is the set of putative motifs $\{m_1, \ldots, m_p\}$, and $\hat{m}^{RS}$ is the best guess at the optimal binding motif $m^*$.

Another approach is to use a regression score (LR score). This is more computationally demanding but can account for some, potentially relevant, confounder information, like di-nucleotide frequencies or sequence length. In order to describe the regression framework, we will rewrite the random set score from Eq. (2). We make the following notations:

$$\mu = \frac{1}{n}\sum_{i=1}^{n} y_i, \qquad \sigma^2 = \frac{1}{n}\sum_{i=1}^{n}(y_i-\mu)^2$$

$$y_i^* = y_i - \mu, \qquad A_j = \frac{n-n_j}{n-1}$$

$$e_j = \frac{1}{n_j}\sum_{i:x_{ij}=1} y_i^*, \qquad \hat{\sigma_j}^2 = \frac{\sigma^2}{n_j}$$

Then we have

$$S^{RS}(m_j) = A_j \times \frac{e_j}{\hat{\sigma_j}} \tag{4}$$

with the top predicted motif $\hat{m}^{RS}$ described by Eq. (3).

Now we can define a linear regression model for the binding interactions, that closely resembles the random set scoring strategy described above. Let the regression coefficient for motif $m_j$ be denoted $\beta_j$, then a simple linear model for binding is:

$$y_i^* = x_{ij}\beta_j + \epsilon_i, \text{with } \epsilon_i \sim N(0, \sigma^2) \tag{5}$$

We estimate the regression coefficient with classical ordinary least squares (OLS): $\beta_j^{OLS} = \frac{1}{n_j}\sum_{i:x_{ij}=1} y_i^*$ and we define the motif enrichment score as follows:

$$S^{LR}(m_j) = \frac{\beta_j^{OLS}}{\hat{\sigma_j}} = \frac{1}{A_j} \times S^{RS}(m_j) \tag{6}$$

The top motif is in this case:

$$\hat{m}^{LR} = \arg\max_{m_j \in M} S_j^{LR} \tag{7}$$

Note that if the size of the motif target set is small relative to the number of all input sequences ($n_j << n$), $A_j \approx 1$ and $S_j^{LR} \approx S_j^{RS}$.

This framework can be extended to account for features of the input sequences that may be unrelated to motif binding. One such potential confounder is the sequence length, and other important features can be derived from the nucleotide content, like the di-nucleotide counts. If we consider $q$ additional confounders, the regression covariates for motif $m_j$ are defined by the following matrix: $Z_j = (x_j, c_1, c_2, \ldots, c_q)$, where $x_j$ represents the column vector of motif matches $x_{ij}$, and $c_k \in \mathbb{R}^n, k \in \{1, \ldots, q\}$ are the confounders. We denote the corresponding regression coefficients with $\beta_{0j}, \beta_{1j}, \ldots \beta_{qj}$, with $\beta_{kj} \in \mathbb{R}, k \in \{0, \ldots, q\}$. Then the model can be expressed with the matrix notation as:

$$Y^* = Z_j\beta_j + \epsilon, \text{where } \epsilon \sim N(0, \sigma^2 I_{n\times n}) \tag{8}$$

In the typical case there are a large number of input sequences and therefore a large number of sample points to use in the estimation process. Also, we need to score a large number of motif candidates, therefore we need a computationally efficient estimator for the regression coefficients, with good statistical properties. We use the simple OLS estimator $\beta_j^{OLS} = (Z_j^T Z_j)^{-1} Z_j^T Y^*$, that provide fast solutions for small to moderate $q$ values. The corresponding motif scoring function is a straightforward generalization of the univariate regression case:

$$S^{LR}(m_j) = \frac{\beta_j^{OLS}}{\hat{\sigma_j}}, \text{with } \hat{\sigma_j}^2 = (\hat{\Sigma}_{\hat{\beta_j}})_{11} \tag{9}$$

where $(\hat{\Sigma}_{\hat{\beta_j}})_{11}$ represents the first diagonal element in the covariance matrix for the parameter estimate $\beta_j^{OLS}$.

We optimized this scoring function in the case of the analyzed CLIP datasets, using as confounders the sequence length and the di-nucleotide counts. For the RNAcompete datasets we optimized the random set scoring function since all the input RNA sequences have the same length and were generated artificially with an uniform model.

#### S1.3 Update rules in the search strategy

Given a motif $m$, a set of candidate motifs is constructed by applying small variations to $m$: in length, sequence, or structure. The $k$-mer $m$ is extended to 16 new $(k+1)$-mers, by independently adding one letter from $A_{basic}$ at one of its end. If $k > 4$, the length of the motif is reduced and 2 new $(k-1)$-mers are considered. Then a large set of new $k$-mers are obtained by changing one letter at a time in terms of structural change or increasing/decreasing sequence degeneracy. Briefly, for each position $j$ in $m$, the following rules are applied:

- if $m[j] \in \{A, C, G, T, a, c, g, t\}$ then four new motif candidates are constructed by replacing $m[j]$, in turns, with its structural complement and the three letters that encode for it and one other nucleotide, keeping the original structure; for example A will become a, M, R, W, and g will become G, k, r, s;

- if $m[j] \in \{W, K, R, Y, S, M, w, k, r, y, s, m\}$ then three new $k$-mers are obtained with $m[j]$ set to either its structural complement, or to one of the nucleotides it encodes, in the same structural context; for example R will become r, A, G;

- if $m[j] \in \{W, K, R, Y, S, M, w, k, r, y, s, m\}$ and $j \notin \{1, k\}$ then a fourth candidate is obtained by setting $m[j] = N$ or $m[j] = n$, depending on the case;

- if $m[j] \in \{N, n\}$ then 10 new motifs are considered by changing $m[j]$ to one letter from either $\{A, C, G, T, W, K, R, Y, S, M\}$, or $\{a, c, g, t, w, k, r, y, s, m\}$.

For example $(4k + 18)$ candidate motifs are generated for a $k$-mer with no sequence degeneracies, or for a $k$-mer with double degeneracy in two middle positions. In the case of a $k$-mer with double degeneracy at its ends, $(4k + 16)$ new motifs are considered.

#### S1.4 Visualization of motif clusters

The k-mer motifs obtained in the search procedure are then clustered in a post-processing step. For a better understanding of this step, **SSMART** plots all unique evolved motifs and the similarity between them in two ways: as a heatmap and as a network graph (see Fig. S3). The similarity is computed with the metric defined for the post-processing step. The heatmap plot depicts all pair-wise similarities between the evolved k-mers, together with their hierarchical clustering. The network graph provides a different perspective of the same data, filtering out the low similarities. In both the heatmap and the network graph the motifs are colored according to the motif cluster they belong to after the custom clustering procedure.

Fig. S3 contains the visualization of k-mers similarity for two libraies corresponding to PUM2 and FUS proteins. While for PUM2 the evolved motifs are more homogeneous, the ones for FUS apear to be more disperse. Depending of the data, there can be small motif clusters (like Motif2 of PUM2 with 4 k-mers or Motif3 of FUS with 5 k-mers), or clusters that incorporate many evolved motifs (like the top PUM2 motif that contains almost half of the k-mers).

#### S1.5 Parameter optimization

While our tool and *Zagros* do not have parameters that need to be set, *RNAcontext* and *GraphProt* have multiple parameters that influence their performance.

*RNAcontext* has three important parameters: the motif length $w$, the structural alphabet $e$ and the number of initializations $s$. The motif length is specified as a range, and *RNAcontext* uses learned models for smaller motifs to initialize longer motif lengths. We set $w$ to $4 - 10$, a range that is consistent with the **SSMART** possible motif lengths. For describing RNA structure, we used the "PHIME" alphabet that consists of five different structures: paired (P), hairpin loop (H), internal loop (I), multiloop (M), and external loop (E). For structure evaluations we considered the paired probabilities. We set parameter $s$ to 3, running the tool with 3 different initializations.

*GraphProt* has six parameters that can be optimized in a dedicated step, using program option $-ls$. For each motif and type of structure used in the synthetic datasets, we ran the optimization procedure on a separate set of sequences and used the optimized values for all datasets in each category. However, the motif recovery rates with the default parameters were better than those obtained with prior parameter optimization (see Table S3), therefore all results reported in the main text correspond to the default values.

#### S1.6 Amount of noise in the synthetic data

In our analyses we generated synthetic datasets that contain specific implanted motifs in various proportions of structured/unstructured binding sites. In addition to the "positive" sequences, we added some noise to each of our 2000 datasets. While for the "primary" synthetic datasets we added in each case 500 noise sequences, as described in the main text, we also generated "secondary" sets with the same positives, but with only 200 noise sequences. When we randomly associated binding scores to sequences in the second datasets, we chose a different distribution, making sure that the 200 noise sequences will be in the bottom 400 scores. Throughtout the paper, the implicit synthetic datasets are the "primary" sets.

The comparison of tool performance on the synthetic datasets with different amount and distribution of noise is presented in Table **??**. For all tools, the recovery rates are better for the case with less noise, but the difference is ussualy bellow 3%. The only exception is *RNAcontext*, with a 15% drop in sequence motif recovery for the "primary" datasets (2000 positives & 500 negatives).

#### S2 Supplementary figures and tables

Table S1: 10 randomly selected PWMs from the RBP compendium. These PWM were used to generate the synthetic datasets used for evaluation. IC refers to information content.

| Motif | Average IC | Consensus | RBP |
|---|---|---|---|
| M159_0.6 | 1.4693 | WGCAUGM | A2BP1, RBFOX2, RBFOX3 |
| M147_0.6 | 1.3348 | GACAGAN | CNOT4 |
| M056_0.6 | 1.253 | ACAACRR | SRSF3 |
| M021_0.6 | 1.2086 | AGGAURA | G3BP2 |
| M232_0.6 | 1.1782 | UUUUUUU | ELAVL1, ELAVL3 |
| M162_0.6 | 1.0972 | AGAAANU | PABPC5 |
| M108_0.6 | 1.0967 | UUUGUUU | ELAVL1, ELAVL3 |
| M242_0.6 | 1.0365 | CCAAAUU | HNRNPR, SYNCRIP |
| M054_0.6 | 0.9501 | GCGCGCG | RBM8A |
| M168_0.6 | 0.6589 | GURGUKU | PSPC1, SFPQ |

Table S2: Biological datasets used in the main text of the manuscript to compare motifs recovered from *in vivo* and *in vitro* data.

| Protein | CLIP SRA accession number | RNAcompete ID |
|---|---|---|
| ELAVL1 | SRR189777 | RNCMPT00032 |
| QKI | SRR048969 | RNCMPT00047 |
| FMR1 | SRR527727 | RNCMPT00016 |
| LIN28A | SRR531465 | RNCMPT00162 |
| LIN28A | SRR458759 | |
| LIN28A | SRR764666 | |
| FUS | SRR070449 | RNCMPT00018 |
| PUM2 | SRR048968 | |
| ROQUIN | SRR857933 | |

Table S3: Comparison of motif recovery rates for *GraphProt* and *Zagros* obtained with default parameters and/or in sequence-only mode. All results correspond to the secondary set of synthetic data, each dataset with 2000 positives and 200 negatives.

| Motif | *GraphProt* | | | *Zagros* | |
|---|---|---|---|---|---|
| | With param. optimization | Default params | Default params & sequence-only | Sequence-structure mode | Sequence-only |
| Sequence | 93.65 | 97.4 | 95.2 | 100 | 96.1 |
| Structure | 72.3 | 74.45 | 20.55 | 19.35 | 24.3 |

Table S4: Comparison of motif recovery rates on two set of synthetic data: the primary one having datasets with 2000 positives and 500 negatives (in a triangular distribution), and the secondary one having datasets with 2000 positives and 200 negatives (situated among the bottom 400 sequences).

| Motif | Synthetic data | SSMART | SSMART-seq | RNAcontext | GraphProt | Zagros |
|---|---|---|---|---|---|---|
| Sequence | 2000 pos & 500 neg (triang) | 91.75 | 92.05 | 58.14 | 94.84 | 100 |
| Sequence | 2000 pos & 200 neg (bottom) | 92.2 | 94.59 | 73.2 | 97.4 | 100 |
| Structure | 2000 pos & 500 neg (triang) | 88.65 | 14.75 | 50.03 | 73.25 | 15.25 |
| Structure | 2000 pos & 200 neg (bottom) | 89.2 | 22.23 | 56.3 | 74.45 | 19.35 |

Table S5: The cutoffs used to distinguish between "recovered" and "not recovered" motifs on two set of synthetic data: (A) for the primary synthetic data (2000 positives and 500 negatives), and (B) the secondary synthetic data (2000 positives and 200 negatives). The tools marked with "-seq" correspond to sequence-only mode, while the "*" marks the *GraphProt* run with optimized parameters.

A)

| Motif finder | Sequence | Structure |
|---|---|---|
| SSMART | 0.8082 | 0.7188 |
| SSMART-seq | 0.8206 | 0.8725 |
| RNAcontext | 0.6995 | 0.7366 |
| GraphProt | 0.6599 | 0.8232 |
| Zagros | 0.817 | 0.8908 |

B)

| Motif finder | Sequence | Structure |
|---|---|---|
| SSMART | 0.8115 | 0.7095 |
| SSMART-seq | 0.8095 | 0.7607 |
| RNAcontext | 0.7008 | 0.7359 |
| GraphProt | 0.6602 | 0.8175 |
| GraphProt* | 0.6514 | 0.8222 |
| GraphProt-seq | 0.5874 | 0.7551 |
| Zagros | 0.818 | 0.8659 |
| Zagros-seq | 0.9092 | 0.7448 |

**A** PUM2 percent of paired bases

| W \ L | 30 | 40 | 50 | 60 | 70 | 80 | 90 | 100 | 110 | 120 | 130 | 140 | 150 |
|---|---|---|---|---|---|---|---|---|---|---|---|---|---|
| L | 0.187 | 0.279 | 0.337 | 0.376 | 0.408 | 0.43 | 0.448 | 0.462 | 0.474 | 0.483 | 0.49 | 0.496 | 0.501 |
| L+10 | 0.243 | 0.313 | 0.359 | 0.394 | 0.418 | 0.438 | 0.453 | 0.466 | 0.476 | 0.483 | 0.491 | 0.496 | 0.506 |
| L+20 | 0.276 | 0.334 | 0.376 | 0.405 | 0.426 | 0.444 | 0.458 | 0.47 | 0.478 | 0.485 | 0.491 | 0.502 | 0.513 |
| L+30 | 0.304 | 0.358 | 0.393 | 0.417 | 0.437 | 0.453 | 0.465 | 0.476 | 0.483 | 0.489 | 0.499 | 0.512 | 0.525 |
| L+40 | 0.321 | 0.374 | 0.405 | 0.428 | 0.446 | 0.46 | 0.47 | 0.481 | 0.488 | 0.496 | 0.509 | 0.523 | 0.532 |
| L+50 | 0.331 | 0.384 | 0.415 | 0.436 | 0.452 | 0.465 | 0.475 | 0.485 | 0.494 | 0.507 | 0.521 | 0.53 | 0.537 |
| L+60 | 0.338 | 0.391 | 0.422 | 0.442 | 0.458 | 0.47 | 0.48 | 0.492 | 0.505 | 0.518 | 0.528 | 0.536 | 0.54 |
| L+70 | 0.344 | 0.397 | 0.427 | 0.447 | 0.462 | 0.474 | 0.487 | 0.502 | 0.516 | 0.526 | 0.533 | 0.538 | 0.542 |
| L+80 | 0.348 | 0.401 | 0.431 | 0.45 | 0.465 | 0.481 | 0.497 | 0.514 | 0.524 | 0.531 | 0.537 | 0.541 | 0.544 |
| L+90 | 0.352 | 0.405 | 0.434 | 0.453 | 0.471 | 0.492 | 0.509 | 0.522 | 0.529 | 0.534 | 0.539 | 0.543 | 0.546 |
| L+100 | 0.355 | 0.408 | 0.436 | 0.459 | 0.482 | 0.503 | 0.517 | 0.527 | 0.533 | 0.537 | 0.541 | 0.544 | 0.548 |

**B** Staufen1 percent of paired bases

| W \ L | 30 | 40 | 50 | 60 | 70 | 80 | 90 | 100 | 110 | 120 | 130 | 140 | 150 |
|---|---|---|---|---|---|---|---|---|---|---|---|---|---|
| L | 0.372 | 0.451 | 0.498 | 0.53 | 0.553 | 0.571 | 0.582 | 0.593 | 0.603 | 0.609 | 0.618 | 0.623 | 0.629 |
| L+10 | 0.412 | 0.473 | 0.511 | 0.539 | 0.56 | 0.573 | 0.585 | 0.596 | 0.604 | 0.612 | 0.619 | 0.624 | 0.632 |
| L+20 | 0.443 | 0.491 | 0.525 | 0.549 | 0.566 | 0.579 | 0.59 | 0.599 | 0.608 | 0.615 | 0.621 | 0.629 | 0.635 |
| L+30 | 0.463 | 0.505 | 0.536 | 0.557 | 0.573 | 0.585 | 0.595 | 0.604 | 0.611 | 0.617 | 0.625 | 0.632 | 0.639 |
| L+40 | 0.474 | 0.516 | 0.544 | 0.563 | 0.578 | 0.59 | 0.599 | 0.606 | 0.613 | 0.621 | 0.628 | 0.637 | 0.642 |
| L+50 | 0.481 | 0.522 | 0.55 | 0.568 | 0.582 | 0.593 | 0.601 | 0.609 | 0.617 | 0.626 | 0.633 | 0.639 | 0.645 |
| L+60 | 0.487 | 0.526 | 0.554 | 0.571 | 0.586 | 0.596 | 0.604 | 0.613 | 0.621 | 0.63 | 0.636 | 0.642 | 0.649 |
| L+70 | 0.491 | 0.529 | 0.557 | 0.574 | 0.588 | 0.598 | 0.609 | 0.617 | 0.626 | 0.633 | 0.638 | 0.645 | 0.652 |
| L+80 | 0.493 | 0.533 | 0.559 | 0.576 | 0.59 | 0.602 | 0.613 | 0.622 | 0.629 | 0.635 | 0.641 | 0.648 | 0.656 |
| L+90 | 0.496 | 0.535 | 0.561 | 0.578 | 0.593 | 0.606 | 0.617 | 0.625 | 0.632 | 0.638 | 0.644 | 0.651 | 0.658 |
| L+100 | 0.498 | 0.536 | 0.563 | 0.581 | 0.597 | 0.61 | 0.619 | 0.628 | 0.634 | 0.641 | 0.647 | 0.653 | 0.661 |

Figure S1: RNAplfold predictions with different values for parameters $W$ (window length) and $L$ (maximum base pair span). The numbers presented in a heatmap-like manner correspond to the percent of paired nucleotides. The values in boxes correspond to three parameter settings widely used in the literature: $(W, L) \in \{(80, 40), (150, 100), (200, 150)\}$. (A) Predictions for PUM2 binding sites. (B) Predictions for Staufen1 binding sites.

Table S6: CLIP datasets used in the main text of the manuscript to compare different motif finders.

| Protein | Protocol | Cell line | Reference | SRR accession number | ID |
|---|---|---|---|---|---|
| ELAVL1 | PAR-CLIP | HEK293 | Kishore *et al.* (2011) | SRR189777 | ELAVL1.1 |
| ELAVL1 | PAR-CLIP | HEK293 | Mukherjee *et al.* (2011) | SRR248532 | ELAVL1.2 |
| ELAVL1 | PAR-CLIP | HeLa | Lebedeva *et al.* (2011) | SRR309285 | ELAVL1.3A |
| ELAVL1 | PAR-CLIP | HeLa | Lebedeva *et al.* (2011) | SRR309286 | ELAVL1.3B |
| PUM2 | PAR-CLIP | HEK293 | Hafner *et al.* (2010) | SRR048967 | PUM2.A |
| PUM2 | PAR-CLIP | HEK293 | Hafner *et al.* (2010) | SRR048968 | PUM2.B |

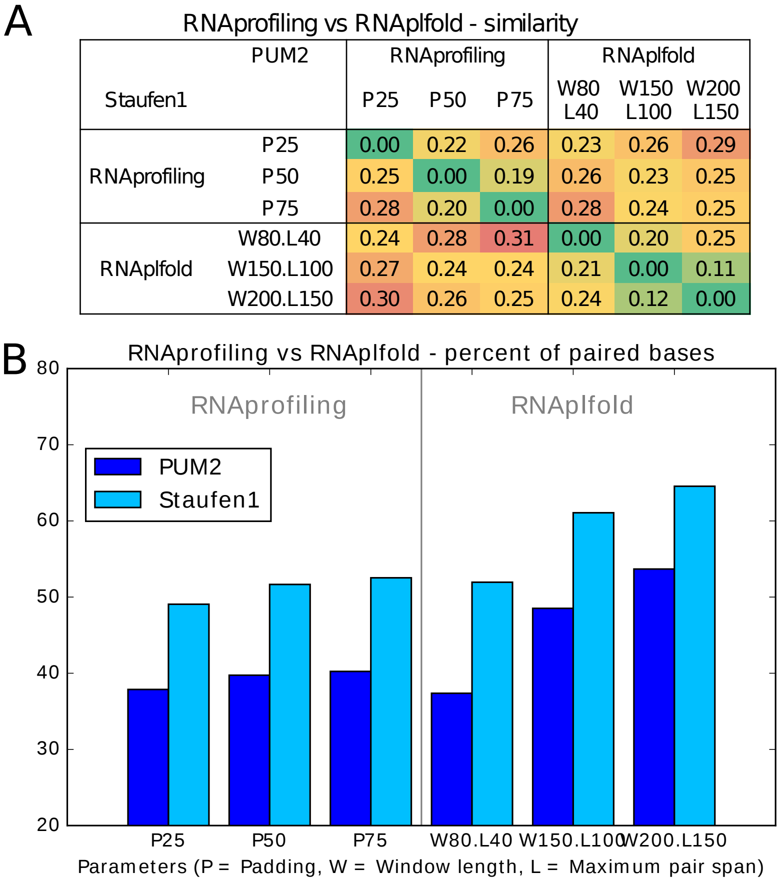


Figure S2: RNAprofiling vs RNAplfold in different parameter settings (P = Padding, W = Window length, L = Maximum pair span). (A) Average similarity between predicted structures. The values above the diagonal correspond to PUM2, while the values below the diagonal correspond to Staufen1. (B) Percent of paired bases for PUM2 and Staufen1 binding sites.

Table S7: P-values obtained with two-sample Kolmogorov-Smirnov tests on the Kendall tau correlation coeficients for the motifs predicted for one specific RBP. The comparison is performed between correlation of the motifs with datasets for the same protein and with datasets for the other considered RBP.

| Tool | ELAVL1 seq motifs | PUM2 seq motifs | ELAVL1 seq-struct motifs | PUM2 seq-struct motifs |
|---|---|---|---|---|
| **SSMART** | 0.0001604 | 0.002797 | 0.0001604 | 0.002797 |
| *GraphProt* | 0.000002719 | 0.3357 | 0.00002447 | 0.002797 |
| *Zagros* | 0.000002719 | 0.06154 | 0.1256 | 0.002797 |

**PUM2_SRR048968 similarities**

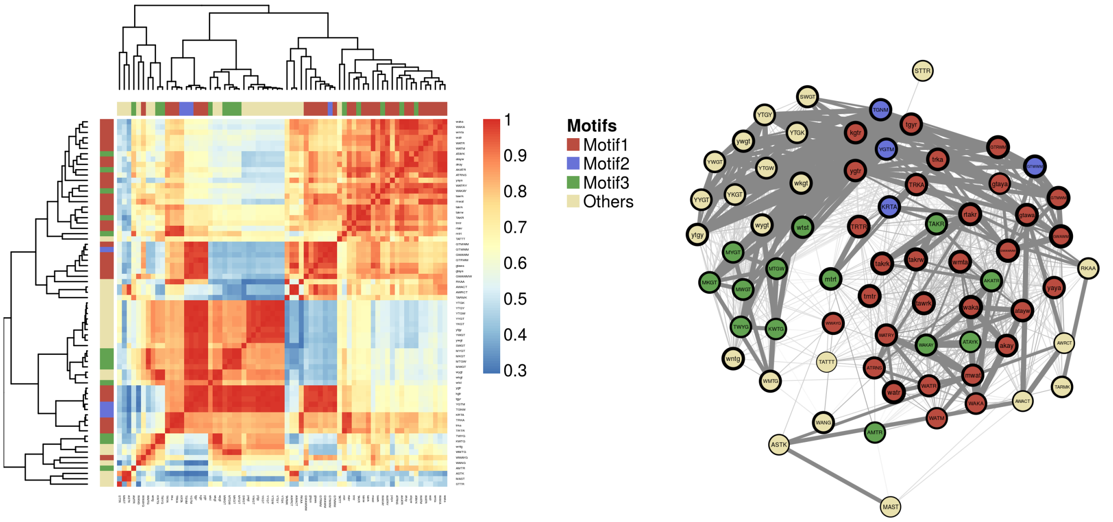


**Heatmap of the similarity scores between all unique evolved k-mers.**
The rows and columns represent k-mers and are color-coded to match the top 3 motifs and all the rest.

**Network graph of all unique evolved k-mers.**
The nodes represent the k-mers (the contour thickness correlates with its SSMART score).
The edges represent the similarity score between k-mers:
thin light gray lines correspond to a score > 0.7, while thick dark grey lines to a score > 0.9

**FUS_SRR070449 similarities**

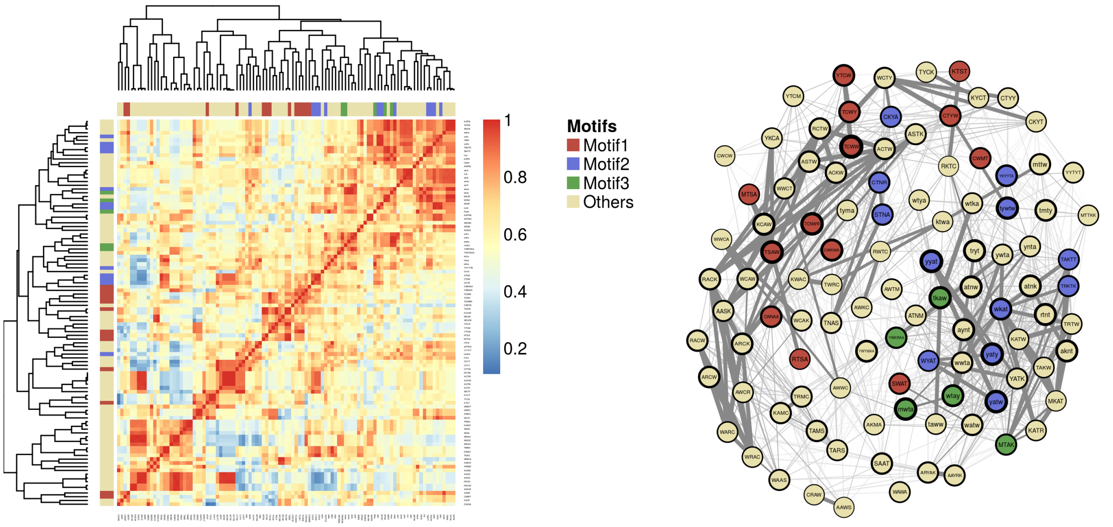


**Heatmap of the similarity scores between all unique evolved k-mers.**
The rows and columns represent k-mers and are color-coded to match the top 3 motifs and all the rest.

**Network graph of all unique evolved k-mers.**
The nodes represent the k-mers (the contour thickness correlates with its SSMART score).
The edges represent the similarity score between k-mers:
thin light gray lines correspond to a score > 0.7, while thick dark grey lines to a score > 0.9

Figure S3: Visualization of motif clusters. Data corresponds to PUM2 (top) and FUS (bottom) proteins. The panel for each RBP contains a heatmap (left) and a network graph (right). Both plots depict the pair-wise similarities between all unique evolved motifs (k-mers). The k-mers are represented on rows and columns in the heatmap plot and as nodes in the network graph. They are color-coded to match the motif cluster they are assign to in the post-processing step. In the network graph, two motifs are connected by an edge if they have more than 90% similarity (thick dark grey edge) or between 70%-90% similarity (thin light grey line). Similarities bellow 70% are not depicted in this plot.

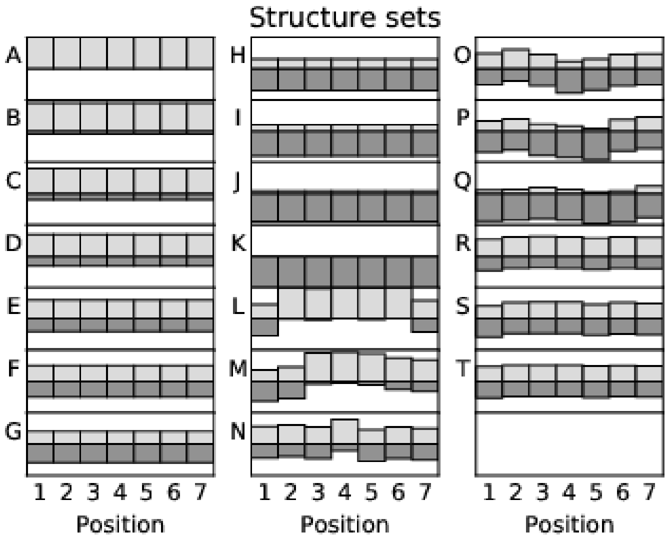


Figure S4: Structural environments in synthetic data. The overall structure that corresponds to the 10 datasets generated for motif M147 is depicted for each type of structure A-T. For each position in the motif, the light grey rectangle coresponds to the probability of being unpaired, while the dark grey represents the paired probability.

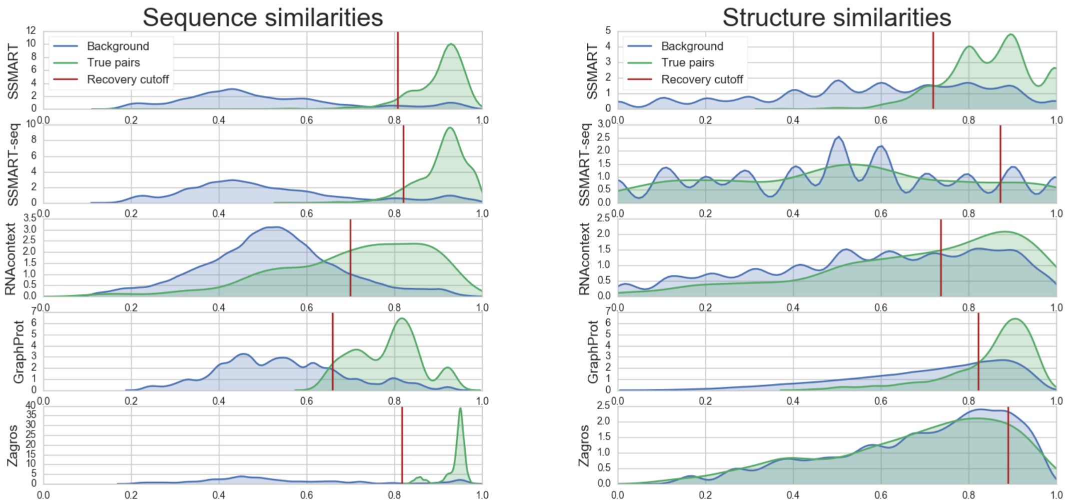


Figure S5: Distribution of similarity scores between predicted and implanted motifs in synthetic datasets. The scores correspond to sequence motifs (left) and structure motifs (right). The similarities computed for all synthetic sets are depicted in green (True pairs), while the background consisting in the similarities between at possible pairs of predicted and implanted motifs are represented in blue. The vertical brown line represents the optimized cuttof for each tool. All motifs that have scores to the rigth of this line are considered "recovered" by the motif finder.

### S3 SSMART results on RNAcompete datasets

The results reported here were obtained using **SSMART** 1.2 with the random set scoring function, since it is fast and there is no need to acount for covariates. All oligos from an RNAcompete experiment have the same length and were designed with a uniform nucleotide distribution.

Table S8: Selected list of RNAcompete datasets on which **SSMART** was applied. They correspond to the selected RBPs for the comparison between motif predictions on *in vivo* and *in vitro* data.

| Protein | RNAcompete ID | GEO ID | Results ID |
|---|---|---|---|
| FMR1 | RNCMPT00016 | GSM1011721 | FMR1 RNCMPT00016 |
| FUS | RNCMPT00018 | GSM1011691 | FUS RNCMPT00018 |
| FXR1 | RNCMPT00161 | GSM1011671 | FXR1 RNCMPT00161 |
| FXR2 | RNCMPT00020 | GSM1011674 | FXR2 RNCMPT00020 |
| IGF2BP2 | RNCMPT00033 | GSM1011697 | IGF2BP2 RNCMPT00033 |
| IGF2BP3 | RNCMPT00172 | GSM1011694 | IGF2BP3 RNCMPT00172 |
| ELAVL1 | RNCMPT00032 | GSM1011629 | ELAVL1 RNCMPT00032 |
| ELAVL1 | RNCMPT00112 | GSM1011563 | ELAVL1 RNCMPT00112 |
| ELAVL1 | RNCMPT00117 | GSM1011582 | ELAVL1 RNCMPT00117 |
| ELAVL1 | RNCMPT00136 | GSM1011621 | ELAVL1 RNCMPT00136 |
| ELAVL1 | RNCMPT00274 | GSM1138954 | ELAVL1 RNCMPT00274 |
| LIN28A | RNCMPT00162 | GSM1011679 | LIN28A RNCMPT00162 |
| QKI | RNCMPT00047 | GSM1011730 | QKI RNCMPT00047 |
| SRSF1 | RNCMPT00106 | GSM1011565 | SF2 RNCMPT00106 |
| SRSF1 | RNCMPT00107 | GSM1011574 | SF2 RNCMPT00107 |
| SRSF1 | RNCMPT00108 | GSM1011594 | SF2 RNCMPT00108 |
| SRSF1 | RNCMPT00109 | GSM1011595 | SF2 RNCMPT00109 |
| SRSF1 | RNCMPT00110 | GSM1011637 | SF2 RNCMPT00110 |
| SRSF7 | RNCMPT00073 | GSM1011738 | SRSF7 RNCMPT00073 |
| SRSF9 | RNCMPT00067 | GSM1011666 | SFRS9 RNCMPT00067 |
| SRSF9 | RNCMPT00074 | GSM1011739 | SRSF9 RNCMPT00074 |

| Protein ID | Reported motif | **SSMART** top 3 sequence-structure motifs |
|---|---|---|
| FMR1 RNCMPT00016 | 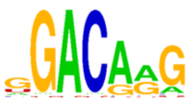 | 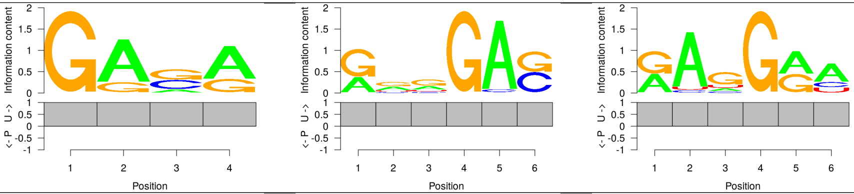 |
| FUS RNCMPT00018 | 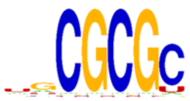 | 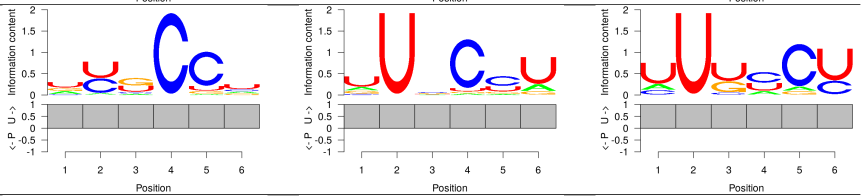 |
| ELAVL1 RNCMPT00032 | 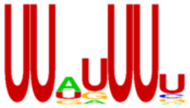 | 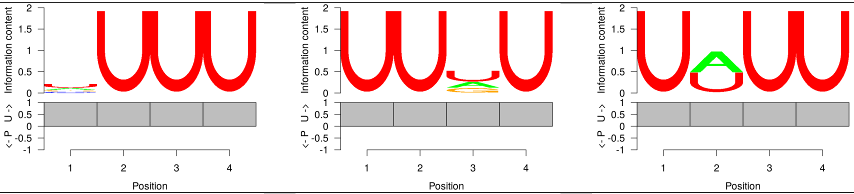 |
| ELAVL1 RNCMPT00112 | 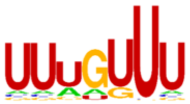 | 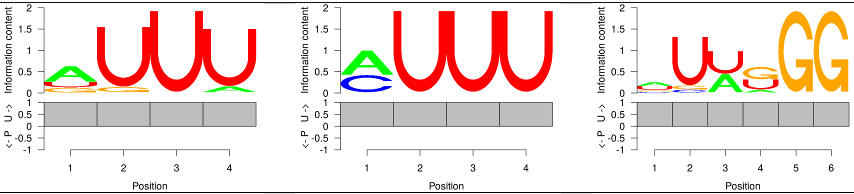 |
| ELAVL1 RNCMPT00117 | 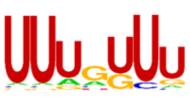 | 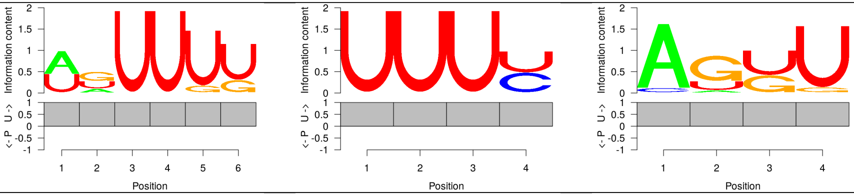 |
| ELAVL1 RNCMPT00136 | 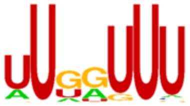 | 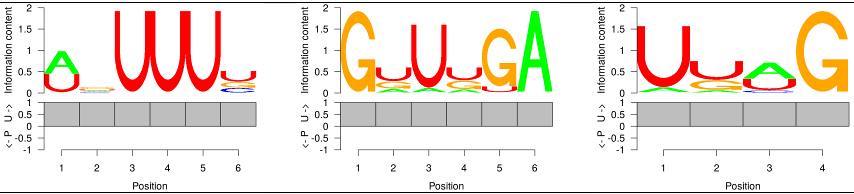 |
| ELAVL1 RNCMPT00274 | 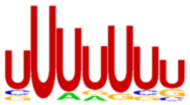 | 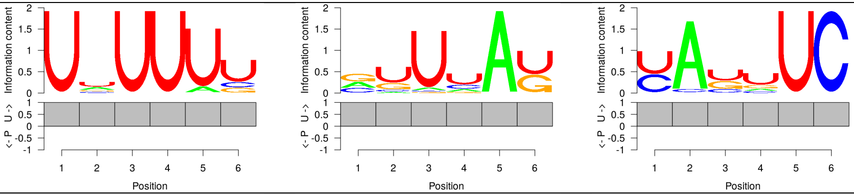 |
| IGF2BP2 RNCMPT00033 | 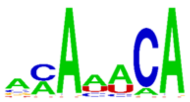 | 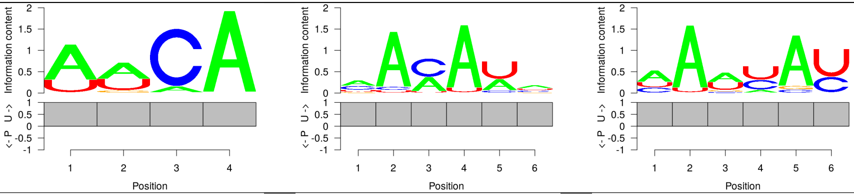 |
| IGF2BP3 RNCMPT00172 | 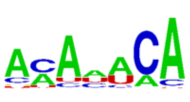 | 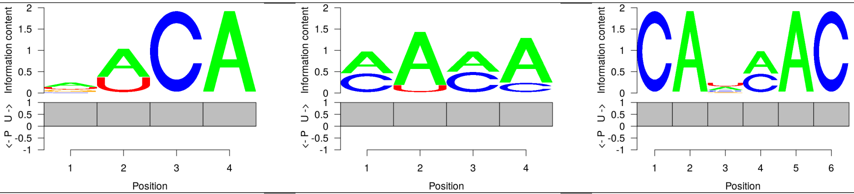 |
| LIN28A RNCMPT00162 | 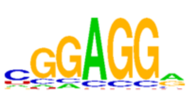 | 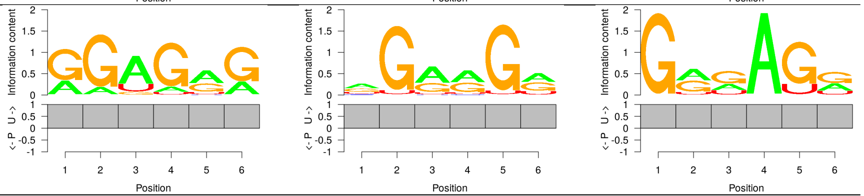 |

| Protein ID | Reported motif | **SSMART** top 3 sequence-structure motifs |
| --- | --- | --- |
| QKI RNCMPT00047 | 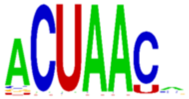 | 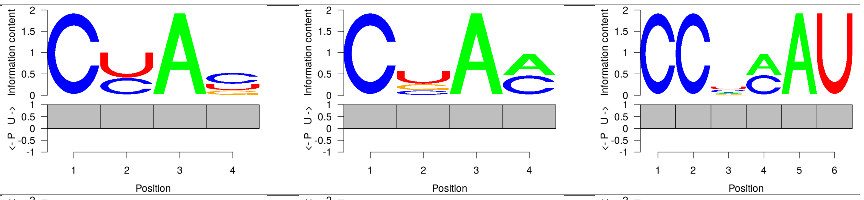 |
| SF2 RNCMPT00106 | 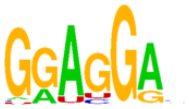 | 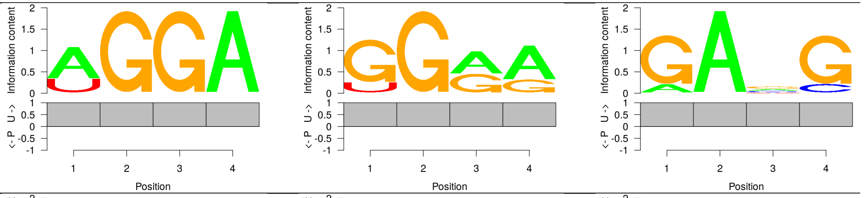 |
| SF2 RNCMPT00107 | 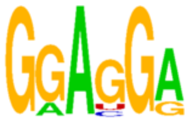 | 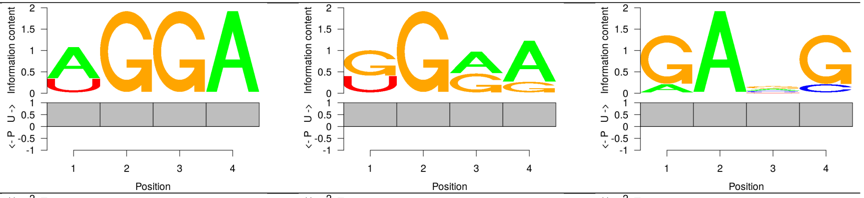 |
| SF2 RNCMPT00108 | 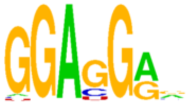 | 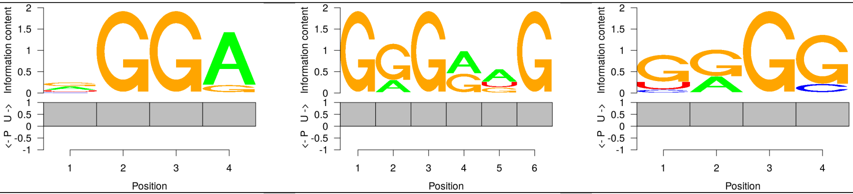 |
| SF2 RNCMPT00109 | 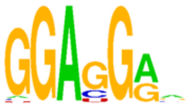 | 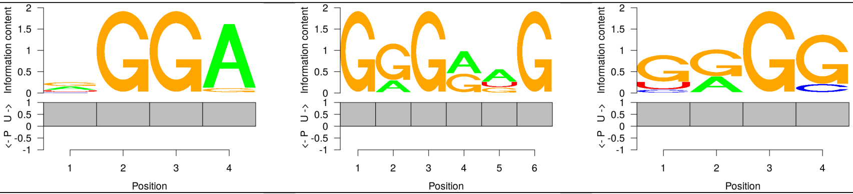 |
| SF2 RNCMPT00110 | 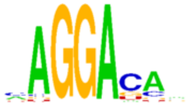 | 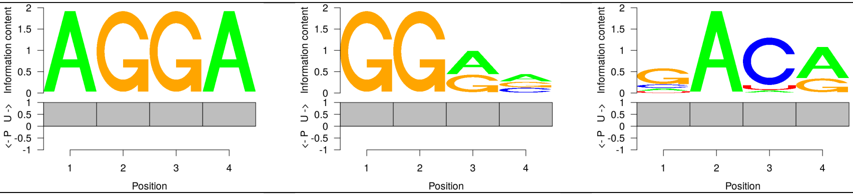 |
| SRSF7 RNCMPT00073 | 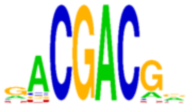 | 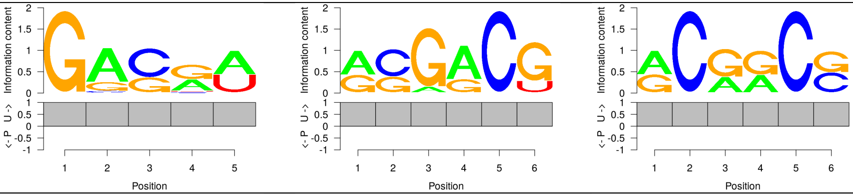 |
| SFRS9 RNCMPT00067 | 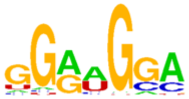 | 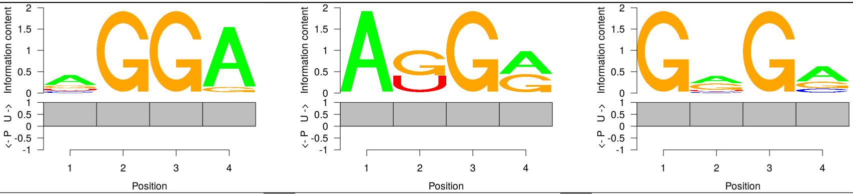 |
| SRSF9 RNCMPT00074 | 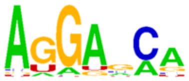 | 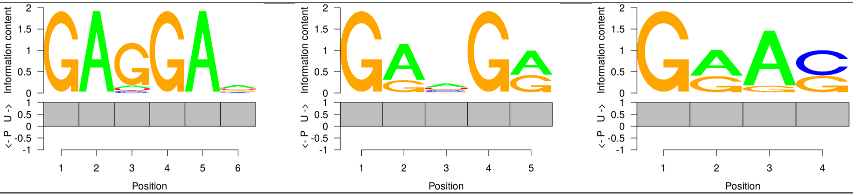 |

### S4 SSMART results on CLIP datasets

The results reported here were obtained using **SSMART** 1.2 with the regression scoring.

Table S9: Full list of CLIP datasets on which **SSMART** was applied.

| Protein | Paper | SRA accession number | Protocol | Cell line | Results ID |
|---|---|---|---|---|---|
| FMR1 | Ascano *et al.* (2012) | SRR527727 | PAR-CLIP | HEK293 | FMR1 SRR527727 |
| FMR1 | Ascano *et al.* (2012) | SRR527728 | PAR-CLIP | HEK293 | FMR1 SRR527728 |
| FUS | Hoell *et al.* (2011) | SRR070449 | PAR-CLIP | HEK293 | FUS SRR070449 |
| FUS | Hoell *et al.* (2011) | SRR070450 | PAR-CLIP | HEK293 | FUS SRR070450 |
| ELAVL1 | Kishore *et al.* (2011) | SRR189777 | PAR-CLIP | HEK293 | ELAVL1 SRR189777 |
| ELAVL1 | Mukherjee *et al.* (2011) | SRR248532 | PAR-CLIP | HEK293 | ELAVL1 SRR248532 |
| ELAVL1 | Lebedeva *et al.* (2011) | SRR309285 | PAR-CLIP | HeLa | ELAVL1 SRR309285 |
| ELAVL1 | Lebedeva *et al.* (2011) | SRR309286 | PAR-CLIP | HeLa | ELAVL1 SRR309286 |
| IGF2BP2 | Hafner *et al.* (2010) | SRR048957 | PAR-CLIP | HEK293 | IGF2BP2 SRR048957 |
| IGF2BP2 | Hafner *et al.* (2010) | SRR048958 | PAR-CLIP | HEK293 | IGF2BP2 SRR048958 |
| IGF2BP2 | Hafner *et al.* (2010) | SRR048959 | PAR-CLIP | HEK293 | IGF2BP2 SRR048959 |
| IGF2BP3 | Hafner *et al.* (2010) | SRR048962 | PAR-CLIP | HEK293 | IGF2BP3 SRR048962 |
| IGF2BP3 | Hafner *et al.* (2010) | SRR048963 | PAR-CLIP | HEK293 | IGF2BP3 SRR048963 |
| IGF2BP3 | Hafner *et al.* (2010) | SRR048964 | PAR-CLIP | HEK293 | IGF2BP3 SRR048964 |
| LIN28A | Cho *et al.* (2012) | SRR458758 | CLIP-seq | A3-1 | LIN28A-mm SRR458758 |
| LIN28A | Cho *et al.* (2012) | SRR458759 | CLIP-seq | A3-1 | LIN28A-mm SRR458759 |
| LIN28A | Cho *et al.* (2012) | SRR458760 | CLIP-seq | A3-1 | LIN28A-mm SRR458760 |
| LIN28A | Wilbert *et al.* (2012) | SRS352780 | CLIP-seq | H9 | LIN28A SRS352780 |
| LIN28A | Wilbert *et al.* (2012) | SRR531465 | CLIP-seq | HEK293 | LIN28A SRR531465 |
| LIN28A | Hafner *et al.* (2013) | SRR764666 | PAR-CLIP | HEK293 | LIN28A SRR764666 |
| LIN28B | Graf *et al.* (2013) | SRR850551 | PAR-CLIP | HEK293 | LIN28B SRR850551 |
| LIN28B | Graf *et al.* (2013) | SRR850552 | PAR-CLIP | HEK293 | LIN28B SRR850552 |
| LIN28B | Hafner *et al.* (2013) | SRR764667 | PAR-CLIP | HEK293 | LIN28B SRR764667 |
| LIN28B | Hafner *et al.* (2013) | SRR764668 | PAR-CLIP | HEK293 | LIN28B SRR764668 |
| LIN28B | Hafner *et al.* (2013) | SRR764669 | PAR-CLIP | HEK293 | LIN28B SRR764669 |
| PUM2 | Hafner *et al.* (2010) | SRR048967 | PAR-CLIP | HEK293 | PUM2 SRR048967 |
| PUM2 | Hafner *et al.* (2010) | SRR048968 | PAR-CLIP | HEK293 | PUM2 SRR048968 |
| QKI | Hafner *et al.* (2010) | SRR048969 | PAR-CLIP | HEK293 | QKI SRR048969 |
| QKI | Hafner *et al.* (2010) | SRR048970 | PAR-CLIP | HEK293 | QKI SRR048970 |
| ROQUIN | Murakawa *et al.* (2015) | SRR857933 | PAR-CLIP | HEK293 | ROQUIN SRR857933 |
| ROQUIN | Murakawa *et al.* (2015) | SRR857934 | PAR-CLIP | HEK293 | ROQUIN SRR857934 |
| SRSF1 | Xiao *et al.* (2016) | SRR2107150 | PAR-CLIP | HeLa | SRSF1 SRR2107150 |
| SRSF3 | Xiao *et al.* (2016) | SRR2107156 | PAR-CLIP | HeLa | SRSF3 SRR2107156 |
| SRSF7 | Xiao *et al.* (2016) | SRR2107152 | PAR-CLIP | HeLa | SRSF7 SRR2107152 |
| SRSF9 | Xiao *et al.* (2016) | SRR2107153 | PAR-CLIP | HeLa | SRSF9 SRR2107153 |
| ZFP36 | Mukherjee *et al.* (2014) | SRR1046759 | PAR-CLIP | HEK293 | ZFP36 SRR1046759 |

| Protein ID | **SSMART** top 3 sequence-structure motifs |
|---|---|
| FMR1 SRR527727 | 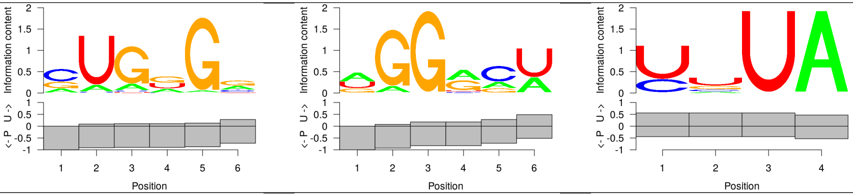 |
| FMR1 SRR527728 | 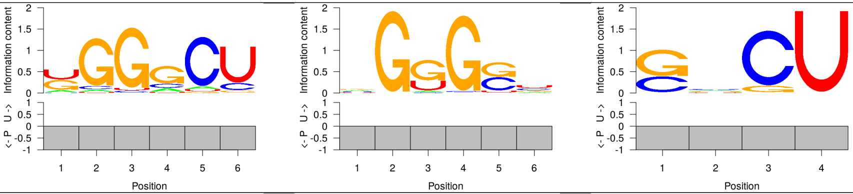 |
| FUS SRR070449 | 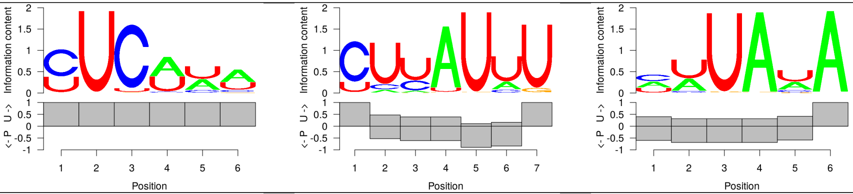 |
| FUS SRR070450 | 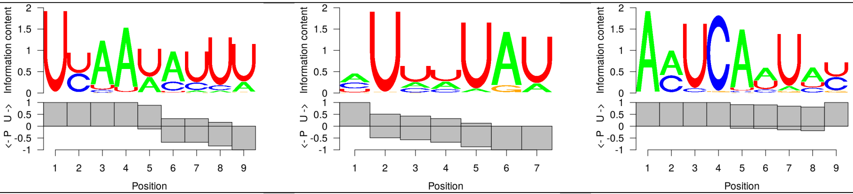 |
| ELAVL1 SRR189777 | 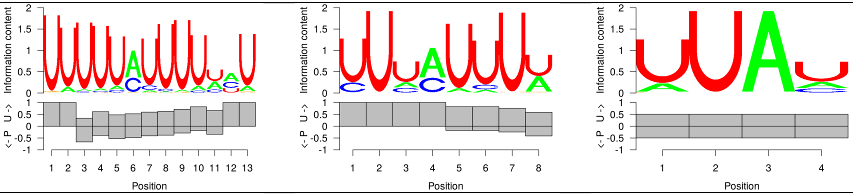 |
| ELAVL1 SRR248532 | 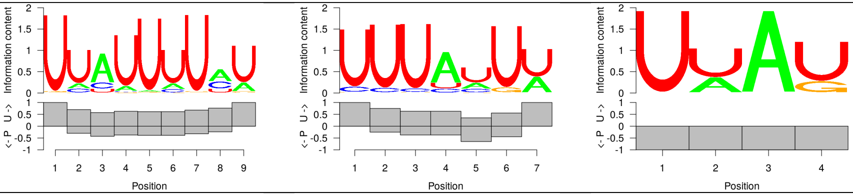 |
| ELAVL1 SRR309285 | 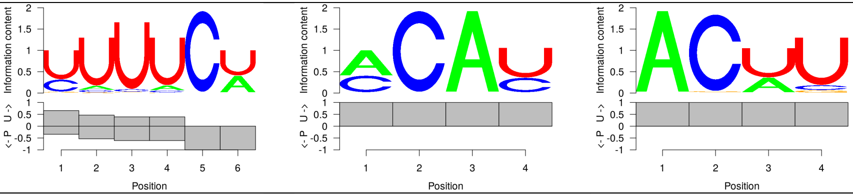 |
| ELAVL1 SRR309286 | 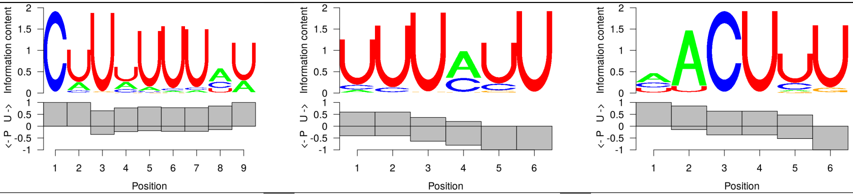 |
| IGF2BP2 SRR048957 | 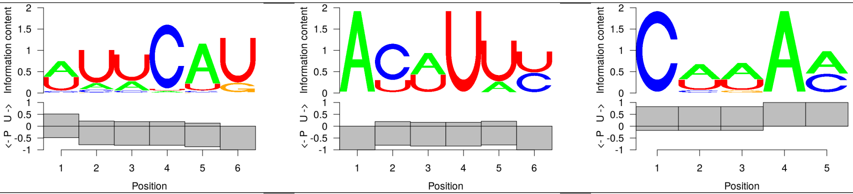 |
| IGF2BP2 SRR048958 | 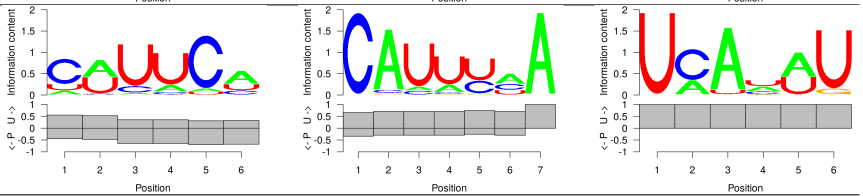 |

| Protein ID | **SSMART** top 3 sequence-structure motifs | | |
|---|---|---|---|
| IGF2BP2 SRR048959 | 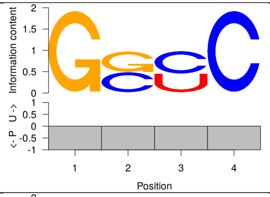 | 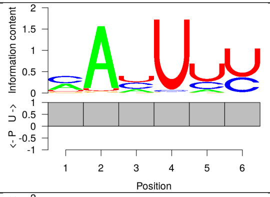 | 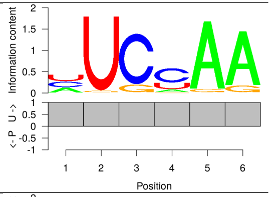 |
| IGF2BP3 SRR048962 | 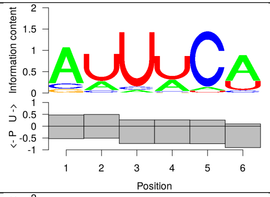 | 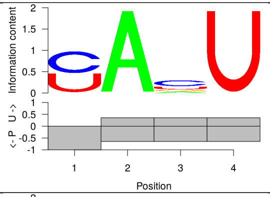 | 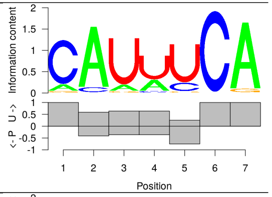 |
| IGF2BP3 SRR048963 | 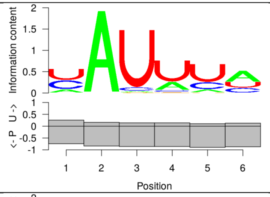 | 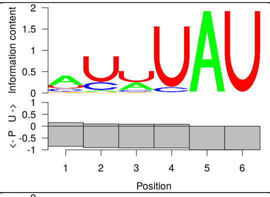 | 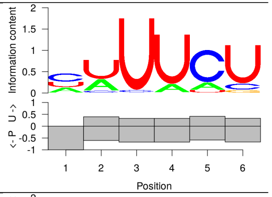 |
| IGF2BP3 SRR048964 | 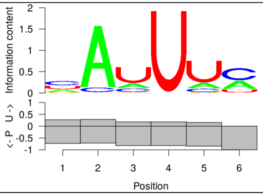 | 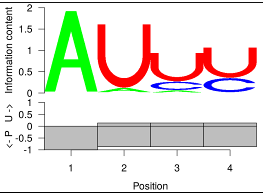 | 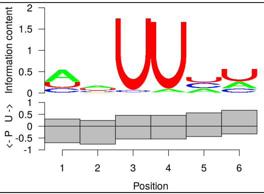 |
| LIN28A-mm SRR458758 | 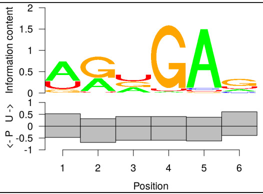 | 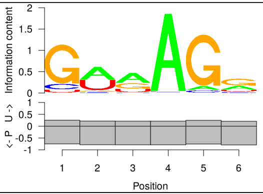 | 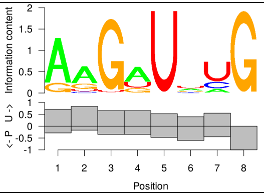 |
| LIN28A-mm SRR458759 | 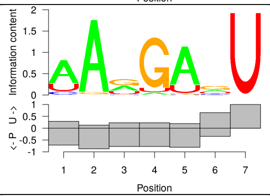 | 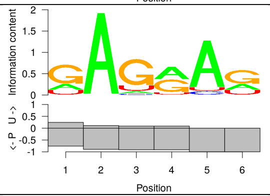 | 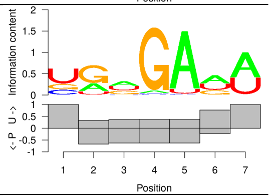 |
| LIN28A-mm SRR458760 | 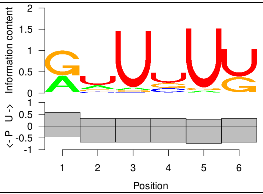 | 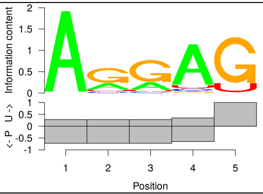 | 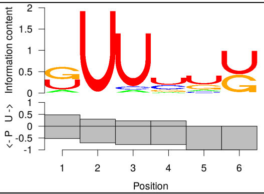 |
| LIN28A SRR531465 | 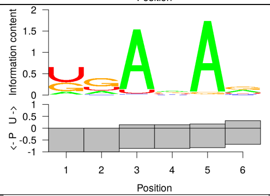 | 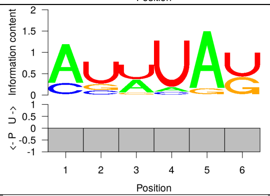 | 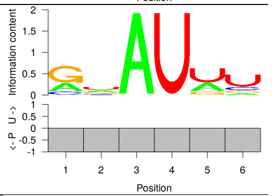 |
| LIN28A SRR531465 | 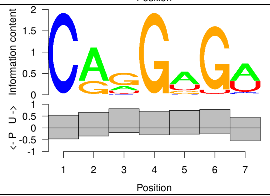 | 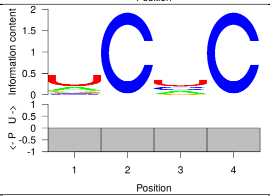 | 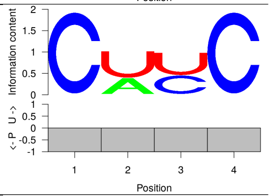 |
| LIN28A SRR764666 | 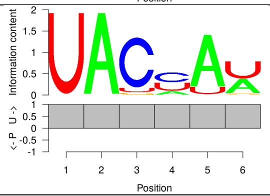 | 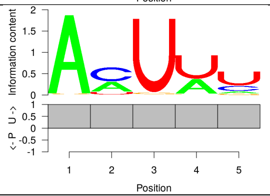 | 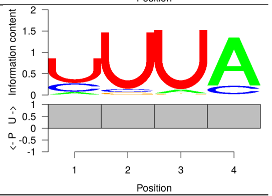 |

| Protein ID | **SSMART** top 3 sequence-structure motifs | | |
|---|---|---|---|
| LIN28B SRR850551 | 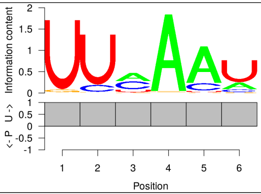 | 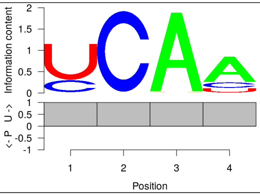 | 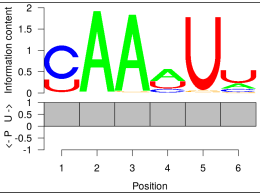 |
| LIN28B SRR850552 | 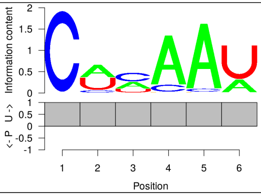 | 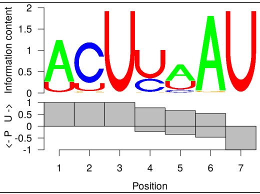 | 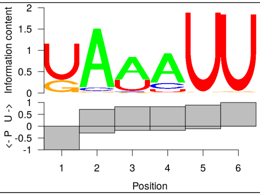 |
| LIN28B SRR764667 | 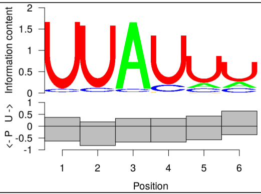 | 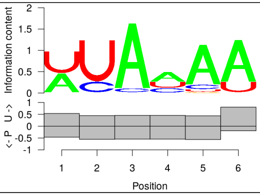 | 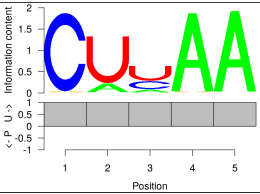 |
| LIN28B SRR764668 | 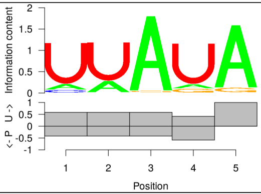 | 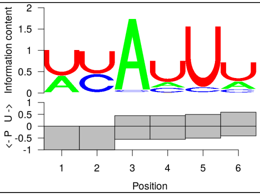 | 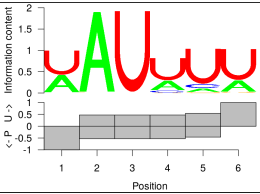 |
| LIN28B SRR764669 | 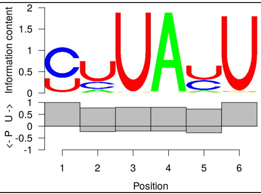 | 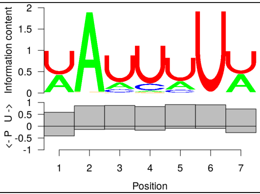 | 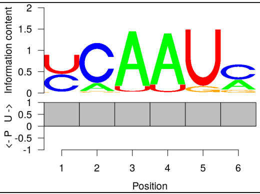 |
| PUM2 SRR048967 | 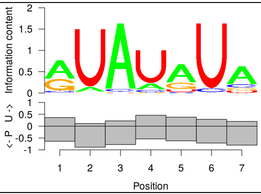 | 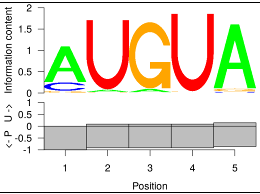 | 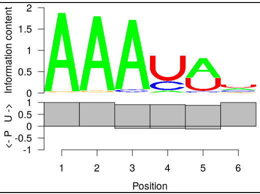 |
| PUM2 SRR048968 | 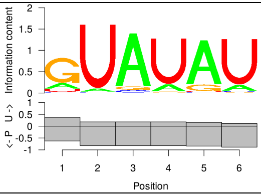 | 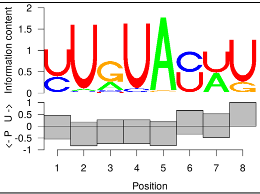 | 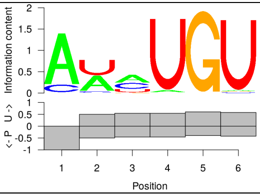 |
| QKI SRR048969 | 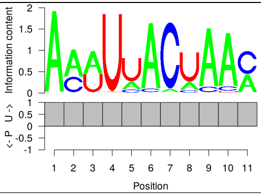 | 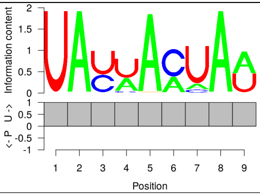 | 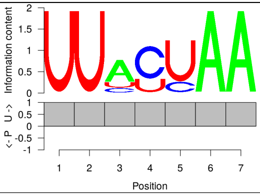 |
| QKI SRR048970 | 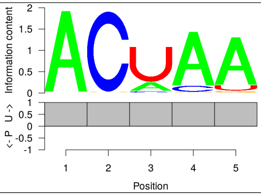 | 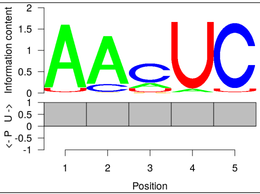 | 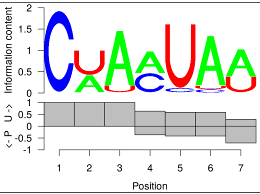 |
| ROQUIN SRR857933 | 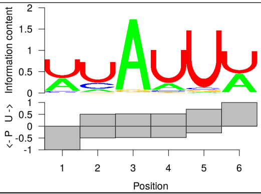 | 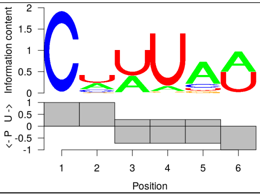 | 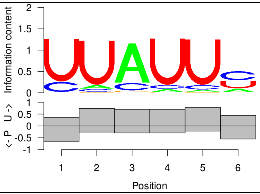 |

| Protein ID | **SSMART** top 3 sequence-structure motifs |
|---|---|
| ROQUIN SRR857934 | 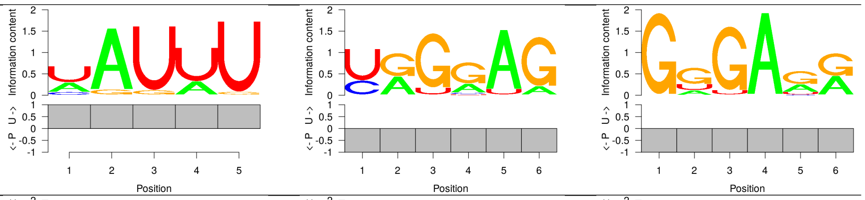 |
| SRSF1 SRR2107150 | 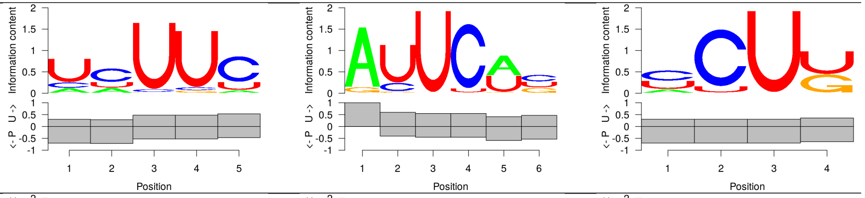 |
| SRSF3 SRR2107156 | 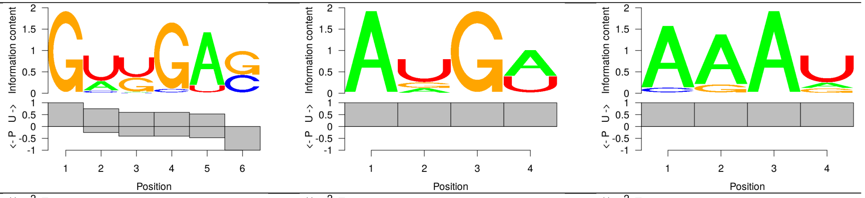 |
| SRSF7 SRR2107152 | 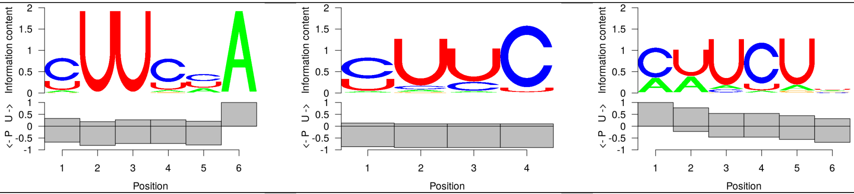 |
| SRSF9 SRR2107153 | 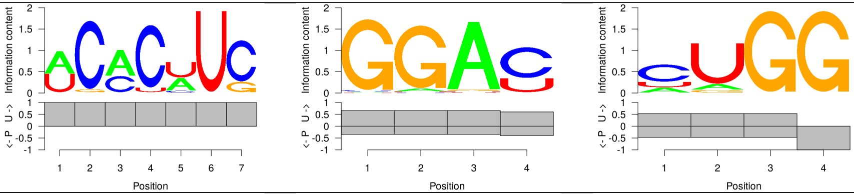 |
| ZFP36 SRR1046759 | 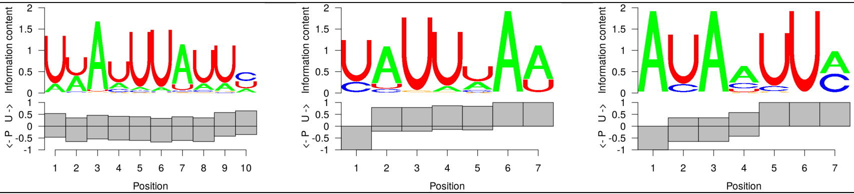 |
